# An iterative gene editing strategy broadens *eIF4E1* genetic diversity in *Solanum lycopersicum* and generates resistance to several potyvirus isolates

**DOI:** 10.1101/2022.07.19.500569

**Authors:** Kyoka Kuroiwa, Benoit Danilo, Laura Perrot, Christina Thenault, Florian Veillet, Fabien Delacote, Philippe Duchateau, Fabien Nogué, Marianne Mazier, Jean-Luc Gallois

**Affiliations:** INRAE, GAFL, F-84143 Montfavet, France; Toulouse Biotechnology Institute, Université de Toulouse, 135 avenue de Rangueil, 31077 Toulouse CEDEX 04, France; INRAE, Agrocampus Ouest, Université de Rennes, IGEPP, F-29260 Ploudaniel, France; CELLECTIS S.A., 8 Rue de la Croix Jarry, 75013 Paris, France; Université Paris-Saclay, INRAE, AgroParisTech, Institut Jean-Pierre Bourgin (IJPB), 78000, Versailles, France.

## Abstract

Resistance to potyviruses in plants have been largely provided by the selection of natural variant alleles of *eukaryotic initiation factors* (*eIF*) *4E* in many crops. However, the sources of such variability for breeding can be limited for certain crop species, while new virus strains continue to emerge. Different methods of mutagenesis have been applied to inactivate the *eIF4E* genes to generate virus resistance, but with limited success due to the physiological importance of translation factors and their redundancy. Here, we employed genome editing approaches at the base level to induce non-synonymous mutations in the e*IF4E1* gene and create genetic diversity in cherry tomato (*Solanum lycopersicum* var. *cerasiforme*). We sequentially edited the genomic sequences coding for two regions of eIF4E1 protein, located around the cap-binding pocket and known to be important for susceptibility to potyviruses. We show that the editing of only one of the two regions, by gene knock-in and base-editing, respectively, is not sufficient to provide resistance. However, combining amino acid mutations in both regions resulted in resistance to multiple potyviruses without affecting its functionality in translation initiation. Meanwhile, we report that extensive base editing in exonic region can alter RNA splicing pattern, resulting in gene knockout. Altogether our work demonstrates that precision editing allows to design plant factors based on the knowledge on evolutionarily selected alleles and enlarge the gene pool to potentially provide advantageous phenotypes such as pathogen resistance.

## Introduction

Recent genome editing techniques allow us to directly inactivate or modify the genome at specific loci, thereby opening the possibility to control specific agronomic traits (Eisenstein, 2022). These techniques, including transcription activator-like effector nucleases (TALEN) and CRISPR (clustered, regularly interspaced, short palindromic repeat)-associated (Cas) endonucleases, allow the targeting of genomic regions of interest and cut the nucleic acid strand(s) to introduce random mutations or to precisely customize target loci with elements like donor sequences and deaminases (Veillet et al., 2020; Gao, 2021). Such precise targeting is essential to minimize the side effects especially for genes that code for proteins involved in several physiological activities through different domains.

Pathogen resistance is a result of evolution of the interaction between plants and pathogens. This is often represented by the natural diversity in resistance alleles of susceptibility genes. These genes can be hijacked by viruses in susceptible plant accessions and are often essential for plant physiology: plants have to develop efficient resistance without impairing their growth. *Eukaryotic initiation factors* (*eIF) 4E* is one of such genes co-evolving with viral-genome linked protein (VPg)-coding gene of potyvirus, the largest plant RNA virus group. *eIF4E*s make a small multi-gene family of mRNA 5’-cap binding proteins that are essential for translation initiation in eukaryotes. The interaction of the potyviruses’ VPg with host eIF4E(s) is essential to establish the infection although the precise mechanism is still unknown (Miras et al., 2017; Tavert-Roudet et al., 2017; Coutinho de Oliveira et al., 2019; Saha and Mäkinen, 2020). Hence, the mutant forms of *eIF4E* have been found as resistant alleles in several crop species including lettuce (*Lactuca sativum*), pea (*Pisum sativum*), and tomato (Ruffel et al., 2002; Nicaise et al., 2003; Gao et al., 2004; Ruffel et al., 2005) though they are not available in all species (Bastet et al., 2017). In pepper (*Capsicum* supp.), 25 allelic forms of *eIF4E1* have been reported and associated with various spectrum and durability of potyvirus resistance (Charron et al., 2008; Moury et al., 2014; Poulicard et al., 2016). Interestingly, those natural mutants harbor only substitutions in their eIF4E1 amino acid sequences compared to their susceptible counterparts. These substitutions are highly clustered in two domains—named Region I and II—of the eIF4E1 protein surrounding the cap-binding pocket (Robaglia and Caranta, 2006; Charron et al., 2008). Despite the importance of this domain for the protein’s original function, many resistance genes have been proven functional reflecting the natural selection of the resulting amino acids (Charron et al., 2008; Moury et al., 2014). This can provide a blueprint to be implemented in resistance breeding.

Similar resistance is found among the ancestral species of tomato. The recessive resistance gene *pot1,* isolated from the wild tomato relative *S. habrochaites* PI247087 accession, encodes an eIF4E1 protein which possesses eight amino acid mutations (L48F, N68K, P69S, A77D, V85L, M109I, K123Q, and N224S) compared to the susceptible cultivated tomato, including five within and in the proximity of Regions I and II. The *pot1*-encoded eIF4E1 protein is associated with a broad resistance spectrum to multiple potyvirus species and strains, while it retains its function to initiate translation (Ruffel et al., 2005; Gauffier et al., 2016).

Various eIF4E knock-out mutants targeting eIF4E1 and/or eIF4E2 have been generated by methods such as TILLING, RNAi, and CRISPR/Cas9 to generate potyvirus resistance in tomato as a model crop. (Piron et al., 2010; Mazier et al., 2011; Gauffier et al., 2016; Schmitt-Keichinger, 2019; Yoon et al., 2020; Kuroiwa et al., 2022). Although the mutant genotypes exhibited resistance to potyviruses, the resistance spectrum associated with single-gene knock-out was limited. Only the severely stunted, double *eif4e1; eif4e2* double knock-out (KO) mutants had a resistance spectrum as large as the one provided by the natural allele *pot1.* This is because most potyviruses can use interchangeably both tomato *Sl*eIF4E1 and *Sl*eIF4E2 (Gauffier et al., 2016; Lebaron et al., 2016). Consequently, KO strategies were achievable, but unlike the natural resistance eIF4Es, neither the single nor double KO of tomato eIF4Es fully met the agronomic demand due to the functional redundancy and the importance of this protein. These results have prompted us to see how more precise genome editing techniques can help designing resistant phenotypes based on the positively selected non-synonymous mutations rather than producing KO alleles in crops like tomatoes (Bastet et al., 2017).

Previously, we provided a proof-of-concept by showing how six mutations selected in the pea (*Pisum sativum*) *eIF4E1*: *sbm1* alleles could be transferred through transgenesis in the *Arabidopsis eIF4E1* susceptible allele and provide resistance to a potyvirus, clover yellow vein virus (Bastet et al., 2018). Moreover, this resistance could be pinpointed to discrete mutations. And when one of those—N176K—was introduced in the susceptible wild-type *Arabidopsis* through base editing, the resulting plants showed the same resistance. This was to our knowledge the first example of using genome precise edition to copy a non-synonymous mutation associated with resistance to viruses (Bastet et al., 2019).

In the present work, we decided to introduce in cultivated tomato a potential amino acid mutation by gene knock-in in the *eIF4E1* Region I, which is known to be important for resistance. Facing the difficulty of such approach, we next decided to randomly tinker another genomic domain in *eIF4E1—*Region II, also known to be involved in resistance, using cytosine base editor (CBE). We show that by iteratively generating diversity in *eIF4E1* genomic regions known to be involved in resistance across plant species, we succeeded in developing a functional potyvirus resistance allele. Incidentally, we observed that heavily mutating a coding region with the CBE can impair the proper gene splicing leading to knock out. Altogether, this work presents the promising potential and the caveat in using genome editing for improvement of crops such as for virus resistance by generating targeted variability in alleles.

## Results

### 1. A codon-change in *SleIF4E1* associated with a P69T mutation is not sufficient to confer resistance to potyviruses

The broad-spectrum resistance allele *eIF4E1*-*pot1* differs from the wild-type susceptible tomato allele by eight non-synonymous codon substitutions, and the role of individual substitutions in the resistance process has not been fully elucidated (Gauffier et al., 2016). Because a related *Solanaceae*, pepper, displays a wide range of resistant alleles, we looked at this *eIF4E1* gene pool to identify the causative mutations for the resistance. In many pepper resistant varieties, one or more mutations at amino acid position 66 to 68, such as P66T, V67E, and A68E within the eIF4E1 Region I, are present (Kang et al., 2005; Charron et al., 2008). As P69 (pepper P66 equivalent) is also mutated in the *pot1*-encoded protein, we decided to introduce a P69T mutation in the susceptible tomato *eIF4E1* allele to generate a new resistant tomato (Supplemental Fig. 1).

**Figure 1.**
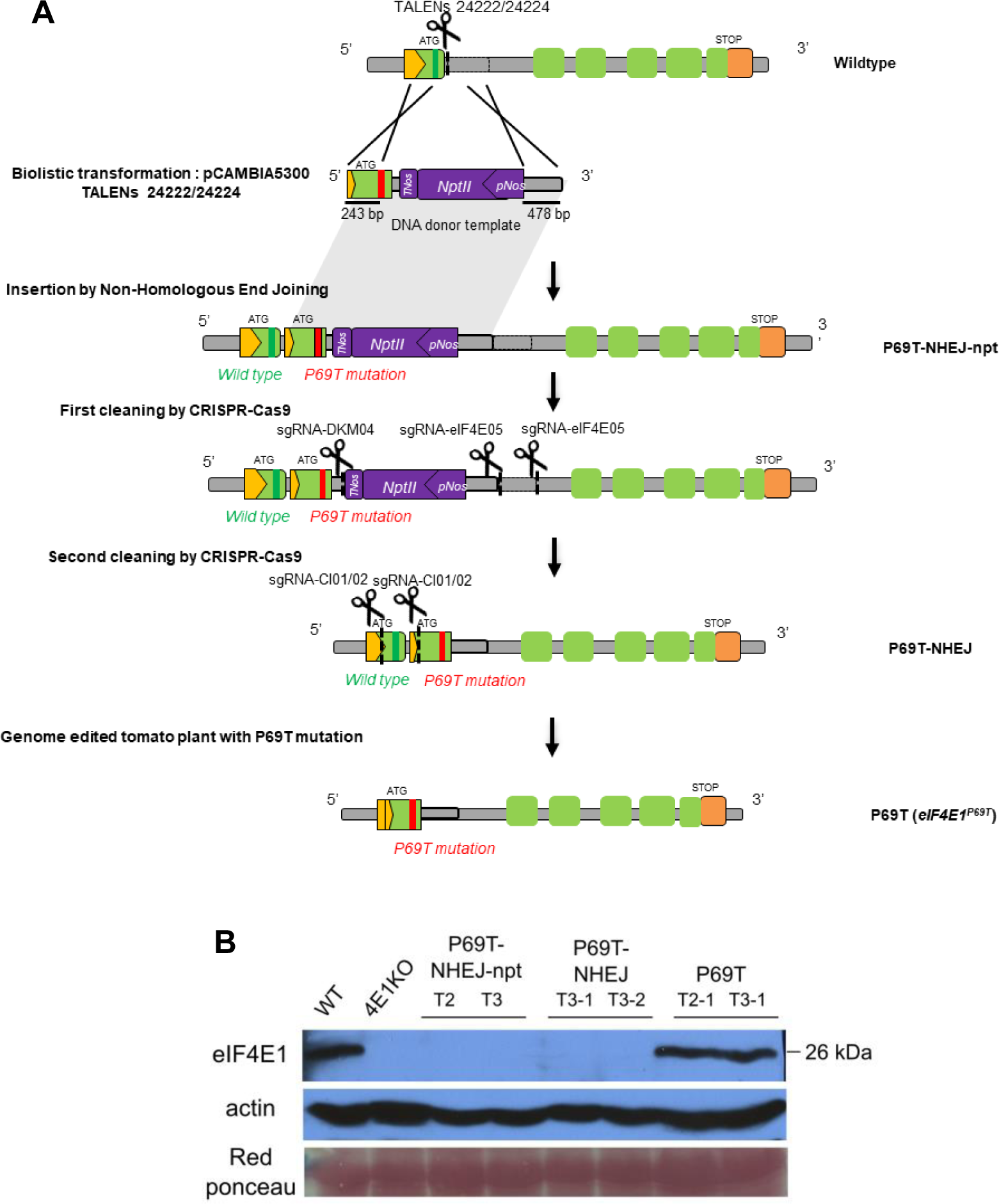
A gene knock-in strategy using TALEN^®^ followed by the targeted elimination of undesirable inserted parts using CRISPR/Cas9 generates a single AA change in *Sl*eIF4E1 exon 1. **A** Schematic diagram of the targeted gene Knock-In and the successive targeted deletion of genomic region. Dotted line under the scissors indicate each targeted site of TALEN or CRISPR sgRNA indicated above. The codon position for P69T mutation is indicated by the strips in green or red (wildtype or mutant). Green boxes represent the exons and yellow pentagon the 5′- UTR. **B** Western blot analysis to detect the *Sl*eIF4E1 accumulation in the homozygous mutant plants from successive genetic modification steps. T2 and T3 indicate the transformant generation. Actin accumulation and red ponceau are shown as the loading control. WT: wildtype, 4E1KO: TILLING *eif4e1^KO^*.

In order to mimic the pepper P66T by mutating codon #69 from CCT to ACA, we attempted a gene knock-in strategy, using a pair of TALENs^®^ to generate a double-strand break in the intron 1 of the *SleIF4E1* gene and to recombine with a donor DNA template. The template contained the *eIF4E1* exon 1 with substituted codon ACA as well as a kanamycin resistance *NptII* gene for selection, flanked by left and right homology arms to promote homologous recombination (HR) (Fig. 1A). Both the TALEN^®^ T-DNA vectors and the donor DNA template were introduced into cells of cherry tomato cultivar (cv.) WVa106 by the biolistic method. Thirty-two regenerating buds were assessed by PCR and sequencing, but none of them harbored an integration of the DNA template via HR (Supplemental Fig. 2). Nevertheless, one plant had integrated the donor template through non-homologous end joining (NHEJ) and named P69T- NHEJ-npt. (Fig. 1A P69T-NHEJ-npt, Supplemental Fig. 2, Supplemental Data 1). This NHEJ-based insertion in the native intron 1 has resulted in the expected insertion of *NptII* gene as well as an undesirable, partial duplication of the *eIF4E1* gene, namely the entire exon 1 deriving from the native and recombined sequences.

**Figure 2.**
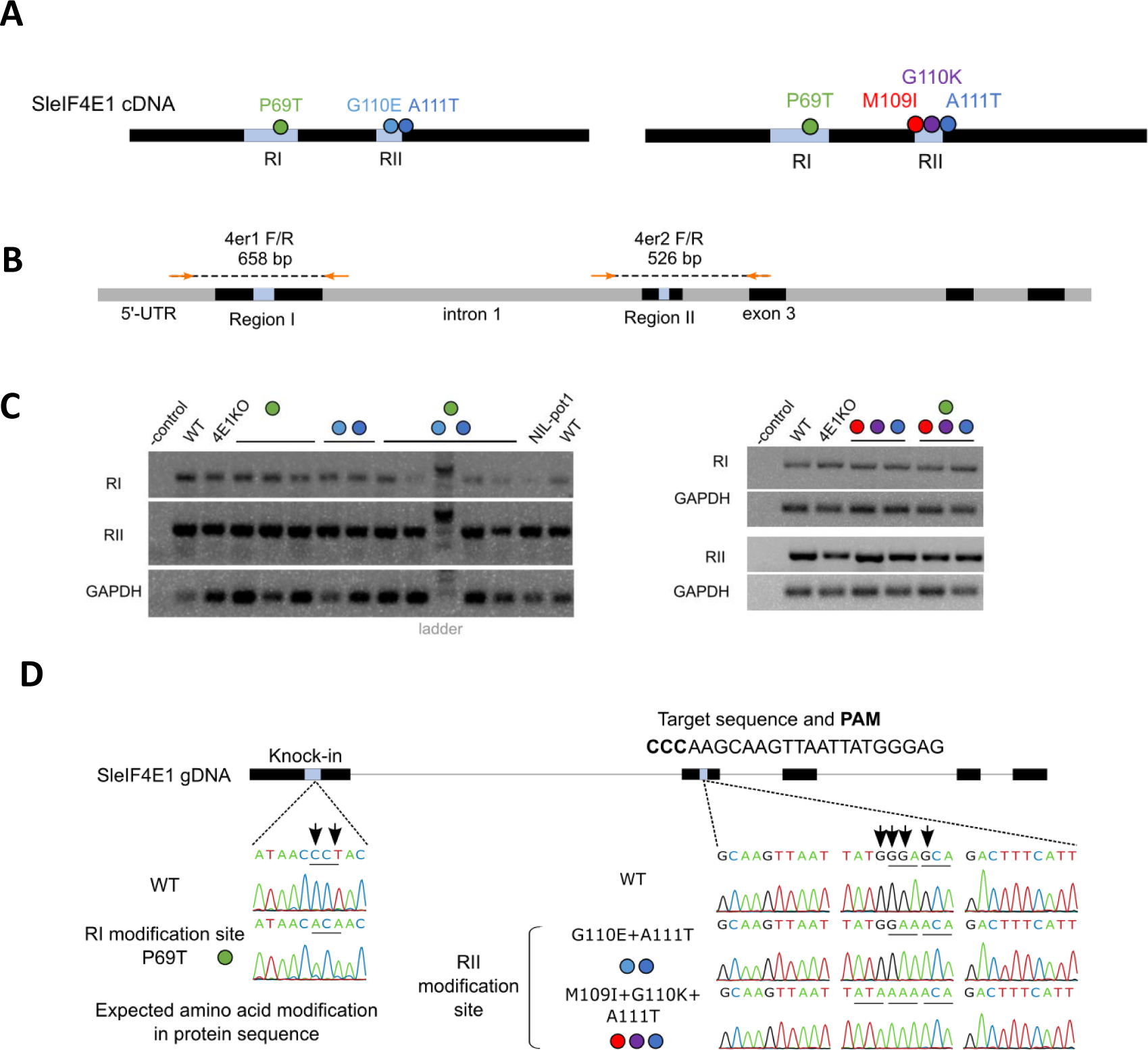
Base-editing in *eIF4E1* Region II generates two combinations of amino acid substitutions in different Region I backgrounds. **A** Schematic diagram of the relative positions of amino acid substitutions induced in this work. Light blue boxes represent Region I and II. **B** Primer positions used in the genomic sequence analyses of *SleIF4E1* Region I and II. Orange arrows are primers. Gray and black box represent introns and exons, respectively. Light blue boxes represent Region I and II. **C** Amplification of genomic sequence of *SleIF4E1* Region I and II. Bands of correct lengths were verified. Two to four independent lines, including those not used in the further analyses, were used per allele. Expected amino acid substitutions from the sequence are represented by the colored circles that correspond to those in diagram A. -control is water. GAPDH is used as internal control. **D** Sanger-sequencing results of the amplified genomic regions confirm the expected modifications in the edited plants. Black boxes indicate the exons in *SleIF4E1* genomic DNA. Representative chromatogram from each modified region (light blue boxes) is shown. Analyses were performed for at least two plants of independent lines of each genotype. Arrows indicate the positions of nucleotide substitutions. The sgRNA in Reverse Complementary sequence used in the base-editing of the Region is shown.

The P69T-NHEJ-npt was selfed to produce TALEN-free plants, homozygous for the modified *eIF4E1* locus. As expected, eIF4E1 protein accumulation was not detected in the progenies using specific polyclonal antibodies: this was consistent with the presence of an approximately 1.4 kb *NptII* expression cassette in the native exon 1 (Fig. 1B, P69T-NHEJ-npt). Thus, in order to restore a functional *eIF4E1* gene, two successive CRISPR/Cas9-mediated editing strategies were used to eliminate undesirable sequences of the recombined *eIF4E1* gene at the 3’ and the 5’ of the insert, respectively (Fig. 1A, alleles P69T-NHEJ and P69T; Supplemental Table 1, 2). Several independent tomato lines were recovered in which both the *NptII* cassette and the endogenous exon 1 of *eIF4E1* were removed. In these plants, the resulting edited *eIF4E1* gene harbored the T69 codon, and the eIF4E1 protein accumulation was restored to wild-type level (Fig. 1B, P69T lanes). These plant lines are hereafter referred to as *eIF4E1^P69T^* lines. This shows that the targeted modification of an exon in *eIF4E1* through knock-in with a DNA donor template is achievable but labor-intensive due to low frequency of homologous recombination and possible rearrangements of the targeted locus.

To evaluate the effect of the eIF4E1 Region I modification on the virus resistance, we used four potyviruses to challenge the plants homozygous at the three editing stages (Fig. 1A) and quantified the viral accumulation by ELISA 21 days after infection. In accordance with the lack of eIF4E1 expression in the first steps of this editing scheme, intermediate lines (P69T- NHEJ-npt and P69T-NHEJ in Fig. 1A and B) were found to be fully resistant to pepper mottle virus (PepMoV) and potato virus Y (PVY) LYE90 isolate but not to other PVY isolates. They therefore present, as expected, the same resistance spectrum as a TILLING *eif4e1^KO^* line (Piron et al., 2010; Gauffier et al., 2016) (Table 1). Interestingly, the *eIF4E1^P69T^*lines in which the eIF4E1 expression was restored were fully susceptible to all four potyvirus isolates tested (Table 1). This implied that introducing a single P69T mutation in the tomato eIF4E1 is not sufficient to induce resistance to the potyvirus infection. However, based on the natural *eIF4E* resistance alleles, which often possess multiple non-synonymous mutations, we reasoned that additional mutations could confer resistance. We thus proceeded to the editing of another genomic region of *eIF4E1*, Region II, where non-synonymous mutations are documented in resistance alleles from multiple species in combination with the Region I.

**Table 1.** Summary of virus infection testing results of each mutant against four potyvirus species.

| Viruses tested | Plant genotypes |  |  |  |  |  |  |
| --- | --- | --- | --- | --- | --- | --- | --- |
|  | WVa106 (wildtype) | TILLING *elf4e1<sup>KO</sup>* | P69T-NHEJ-npt T3-1 | P69T-NHEJ T3 |  | P69T |  |
|  |  |  |  | 1 | 2 | 1 | 2 |
| PepMoV | S | R | R | R | R | S | S |
| PVY-LYE90 | S | PR | PR/S | PR | PR | S | S |
| PVY-N605 | S | S | S | S | S | S | S |
| PVY-SON41 | S | S | S | S | S | S | S |

The accumulation of viral coat protein (CP) 21 days post inoculation (dpi) was measured by ELISA. Six plants are used for each mutant line represented by column (two lines for the P69T-NHEJ and P69T genotypes) and 12 for wildtype. S: susceptible, R: resistant (colored in yellow), PR: partially resistant (CP level lower than S but higher than R; colored in orange).

### 2. A base editing strategy efficiently generates diverse alleles with multiple substitutions in *eIF4E1* Region II

Since many natural potyvirus resistance alleles possess amino acid substitutions in both Region I and II of eIF4E1, we decided to target the Region II genomic sequence in *eIF4E1^P69T^*plants. The same editing strategy was carried out in parallel on wild-type WVa106 plants. In order to bypass the labor-intensive knock-in strategy, we chose a base-editing strategy using nCas9-cytidine deaminase enzyme (or cytosine base editor; CBE). Because a CBE can introduce mutations at one or more cytosines, provided they are located at appropriate position related to a PAM sequence (Nishida et al., 2016), it can trigger varied amino acid changes in this region instrumental for potyvirus resistance. An *SleIF4E1* guide RNA was designed to modify a maximum of four cytosines situated in the editing window at 16-20-nt relative to the PAM on the reverse strand, with potential to induce amino acid changes for M109, G110 and A111 (Supplemental Fig. 3A). Moreover, to facilitate the selection, we implemented a co-editing strategy which can simultaneously mutate cytosines in the *acetolactate synthase SlALS1* targeted genomic regions (Supplemental Fig. 3C). As shown previously, plants edited for *SlALS1* can be selected on ALS inhibitors such as chlorsulfuron, allowing a selection of potentially edited plantlets by CRISPR activity. (Veillet et al., 2019).

**Figure 3.**
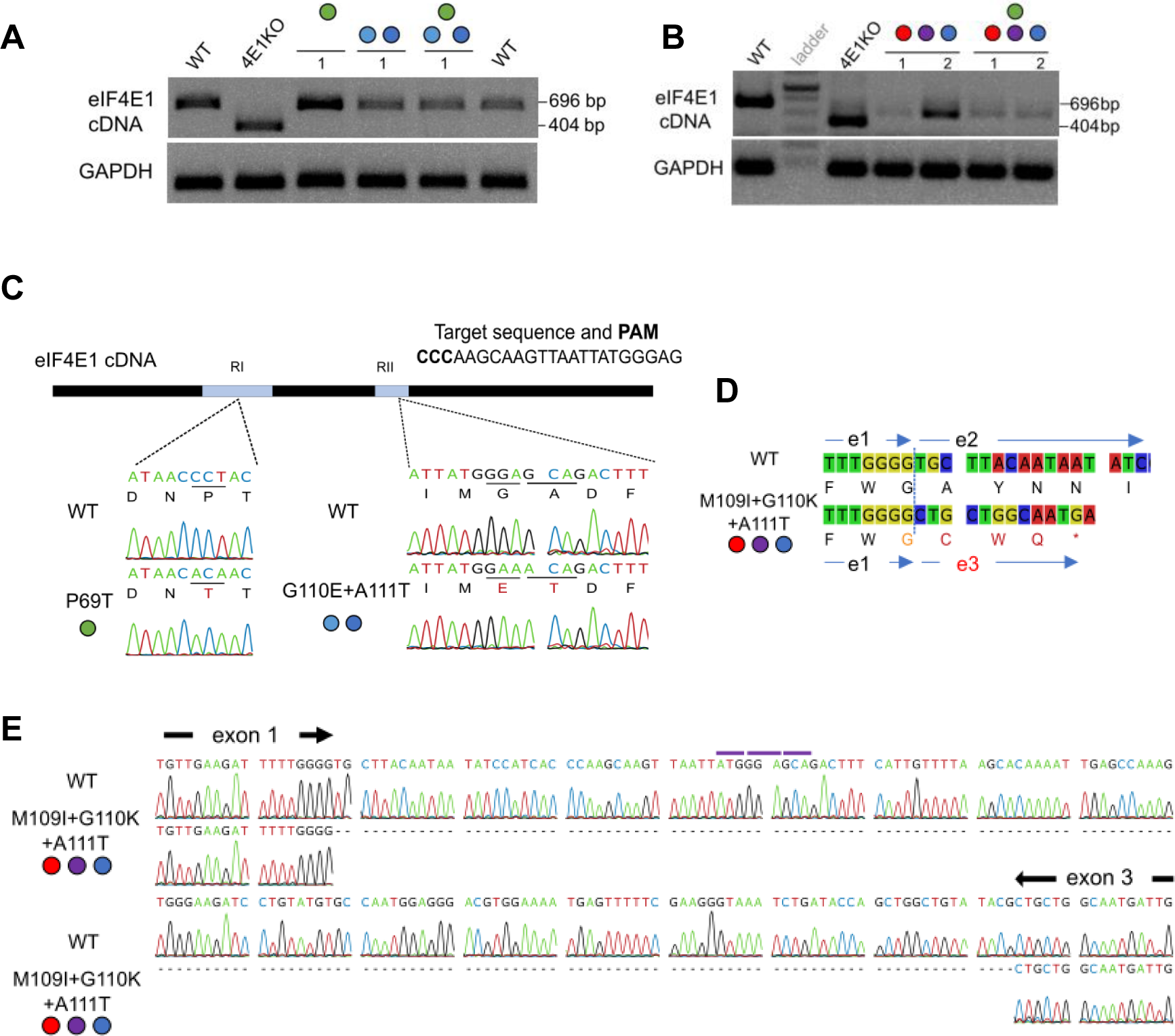
*eIF4E1* cDNA analysis confirms correct mRNA expression for moderately edited plants but reveals abnormal splicing for the triple-codon mutants in Region II. **A, B** Amplification of *eIF4E1* cDNA following reverse transcription; the expected size of the PCR fragment is 696 bp for WT and 404 bp for the TILLING *eif4e1^KO^*. The GAPDH cDNA is amplified as internal control. Representative sample from two independent mutant lines are shown. **C** Sanger-sequencing of the amplified *SleIF4E1* cDNA confirms the expected modifications in the P69T- and G110E+A111T-coding mutant lines. Representative chromatogram from each modified region is shown. Analyses were performed for at least two plants of independent lines of each genotype. Expected amino acid substitutions from the genomic DNA sequencing are represented by the colored circles (cf. Fig. 2A). Predicted translation of each sequence is shown under the nucleotide sequence. Amino acid substitution relative to WT sequence is indicated in red. **D** Predicted protein translation of cDNAs from the WT plant and from an IKT-containing edited plant. The premature stop codon in the eIF4E1 cDNA from IKT plants is represented by a star. e1, e2, and e3 are exons 1, 2, and 3. **E** Chromatogram of the representative cDNA sequence of IKT group shows an unexpected deletion corresponding to the wild-type exon 2. Shown from the 3′ end of the exon 1 to the 5′ end of the exon 3. Purple bars indicate the positions of the codons expected for mutations based on the genomic DNA sequencing.

Explants that grew on the chlorsulfuron selection media were recovered from both WVa106 and *eIF4E1^P69T^* backgrounds. Based on the sequencing of the genomic DNA covering *eIF4E1* exon 2 that includes the targeted Region II, two edited profiles were selected 1) a double amino acid change of G110E + A111T (hereafter referred to as ET) and 2) a triple change of M109I + G110K + A111T (hereafter referred to as IKT) (Fig. 2A). We therefore obtained four original combinations of mutations in eIF4E1 that have never been described among the natural diversity of eIF4Es. Moreover, this setup allows to potentially monitor the effect of mutations in the two Regions, independently or together. Two independent lines for each of the four new alleles (*eIF4E1^ET^*, *eIF4E1^P69T/ET^*, *eIF4E1^IKT^*, and *eIF4E1^P69T/IKT^*) were selected and after three selfings, transgene-free homozygous lines for the *eIF4E1* modification were obtained (Fig. 2B- D).

### 3. Extensive base editing in exon 2 is associated with mis-splicing of *eIF4E1* mRNA

To characterize the effects of modifications on *eIF4E1* gene function in each mutant line, we performed RT-PCR on the *eIF4E1* mRNA using specific primers. In the *eIF4E1^P69T^*, *eIF4E1^ET^*, and *eIF4E1^P69T/ET^*-edited lines, cDNA of the same size as wildtype (696 bp), was amplified (Fig. 3A). A shorter cDNA of 404 bp was amplified for *eif4e1^KO^*mutant which is a product of an aberrant splicing due to a mutation in a 5′-splice site flanking the exon 3 (Piron et al., 2010). In contrast, in the *eIF4E1^IKT^*and *eIF4E1^P69T/KT^* lines, cDNA with a size smaller than wildtype but larger than that of *eif4e1^KO^*mutant was amplified (Fig. 3B). Based on the Sanger-sequencing and alignment with the wild-type sequence, all but the *eIF4E1* cDNA derived from *eIF4E1^IKT^*and *eIF4E1^P69T/KT^* contained the predicted sequences including the correct nucleotide substitutions already verified in genomic DNA (Fig. 2D, 3C). In contrast, the shorter cDNAs amplified from all of the IKT modification mutants were missing a 166-bp-long segment which entails Region II of *eIF4E1*, rendering the cDNA shorter (530 bp-long) (Fig. 3E). In fact, this missing segment perfectly matched the entire exon 2.

To gain insight into this event, we analyzed closely the amplified gDNA sequences surrounding the Region I and II and the aberrant cDNA sequence obtained from the edited tomato lines in comparison with previous studies on regulatory elements. In the gDNA fragment that spans the 526-bp genomic region from the last 100 bp of the intron 1 to the first 30 bp of the exon 3 (amplified by 4e2F/R in Fig. 2B), we did not observe any other mutations than our predicted target codon modifications that code for the IKT substitutions. The sequencing also revealed that the exon 1 sequence was followed by GT and the exon 3 preceded by AG in the wild-type and edited genome (Supplemental Fig. 4A). These correspond to the highly conserved dinucleotides known as the donor and acceptor sites for the pre-mRNA splicing (Brown and Simpson, 1998; Li et al., 2019). They are indeed the donor site and the acceptor site of the intron 1 and 2 in the wild-type *SleIF4E1*, respectively (Piron et al., 2010). The splice sites flanking the exon 2 sequence were also intact in the analyzed gDNA (Supplemental Fig. 4A). Thus, we concluded that the modification of a wild-type sequence ATGGGAGCA into an IKT-coding sequence ATAAAACA in the exonic region has altered the splicing pattern and induced an exon skipping in *eIF4E1* (Supplemental Fig. 4B).

**Figure 4.**
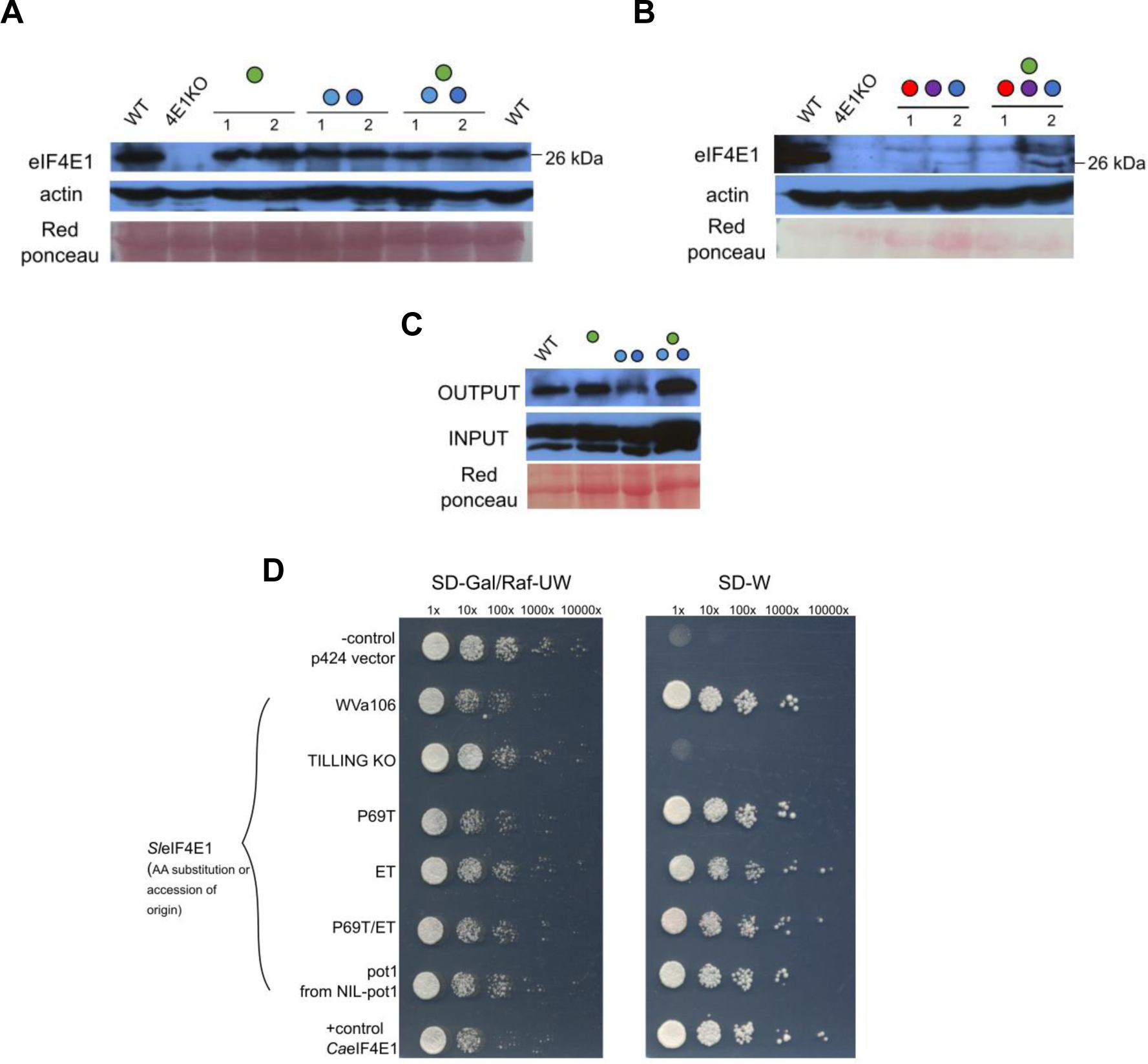
Moderate editing in eIF4E1 region II does not affect the protein accumulation level *in planta* and the protein functionality *in vitro* and *in vivo*. **A, B** Western blot analysis to detect the *Sl*eIF4E1 accumulation in the edited lines. Actin accumulation and red ponceau are shown as the loading control. Genotypes are indicated by the colored circles (cf. Fig. 2A). Two independent edited lines were used per genotype. The expected molecular weight of *Sl*eIF4E1 is 26 kDa. **C** Cap-affinity purification and following western blot shows the cap-binding ability of the edited eIF4E1 proteins at wild-type level. OUTPUT is immunoblot using anti-SleIF4E1 specific antibody on the eluted samples after incubation with m^7^GTP-cap analog and repeated washing. INPUT is immunoblot using anti-actin antibody on the total soluble protein extracts as the loading control. Red ponceau staining of the INPUT loading is also shown as the loading control. **D** Yeast complementation assay demonstrates that the edited *SleIF4E1* proteins correctly expressed *in planta* retain the functionality in translation initiation. Gal/Raf-UW is the control media and Glu-W is the selective media for the eIF4E function. J055 transformed with p424 empty vector is shown as a negative control. *Ca*eIF4E1 is the positive control. All *Sl*eIF4E1 are the transformants with the cDNA obtained from the edited plants. X indicates the dilution level of the yeast culture. Ca: *Capsicum annuum*

At translational level, this exon deletion would cause a frameshift at position 97 in amino acid sequence, resulting in a premature termination codon corresponding to position 100 in the wild-type sequence (Fig. 3D). This possibly leads to truncated eIF4E1 production and/or the degradation of such aberrant form in *eIF4E1^IKT^* and *eIF4E1^P69T/IKT^* lines.

### 4. Mis-splicing resulting from heavy editing at *eIF4E1* Region II affects the eIF4E1 protein accumulation while a moderate editing maintains accumulation and functionality of eIF4E1

Not only the mis-splicing, but also the introduction of amino acid modifications itself can destabilize the protein structure affecting the accumulation and/or functionality of the protein. To verify the protein accumulation, we performed western blot analysis on the proteins extracted from leaves for each edited line. All independent lines of plants with the *eIF4E1^P69T^*, *eIF4E1^ET^*, and *eIF4E1^P69T/ET^* alleles expressed eIF4E1 protein (26 kDa) at a level comparable to that of the wildtype (Fig. 4A). In contrast, lines harboring three mutations in eIF4E1 Region II (IKT) showed no to very low accumulation of eIF4E1 at the expected size (Fig. 4B). This suggests that the amino acid substitutions in the eIF4E1^P69T^, eIF4E1^ET^, and eIF4E1^P69T/ET^ do not affect the protein accumulation. The results also implied that the mis-splicing due to the IKT modification in Region II has led to lack of accumulation or expression of the eIF4E1 protein. As a consequence, the *eIF4E1^IKT^* and *eIF4E1^P69T/IKT^* alleles could be considered *eIF4E1* loss-of-function alleles.

Given the importance of eIF4E1 protein in plant translation, we further checked, in addition to the protein accumulation, whether the functionality of the edited eIF4E1 proteins was not affected. To verify the ability of the edited proteins to bind the 5′-cap structure and initiate translation, we performed both *in vitro* and *in vivo* methods. We first assessed the ability of the eIF4E1 proteins to bind the m^7^GTP-cap analog. After the incubation of total protein extract with cap analog, eIF4E1 protein was detected for the wildtype as well as for all three edited lines, namely *eIF4E1^P69T^*, *eIF4E1^ET^*, and, *eIF4E1^P69T/ET^*, showing that the edited alleles code for eIF4E1 with cap-binding ability (Fig. 4C). Additionally, we used yeast complementation assay, which relies on the conditional complementation of a yeast strain lacking its native *eIF4E* gene. In accordance with the cap affinity purification results, the heterologous expression of the three edited tomato alleles complemented the lack of a native yeast *eIF4E* (Fig. 4D). The serial dilution suggests that the complementation efficiency with the edited version of eIF4E1 is comparable to that achieved with the wild-type eIF4E1 as well as the *pot1*-encoded eIF4E1 (Fig. 4D). Therefore, the *in vitro* and *in vivo* assays strongly suggest that the editing carried out in *eIF4E1^P69T^, eIF4E1^ET^,* and *eIF4E1^P69T/ET^* alleles did not affect their functionality in translation initiation processes, in a similar fashion to the natural *eIF4E* varieties.

Taken together, these results show that the P69T editing in Region I does not affect eIF4E1 expression or function. In contrast, while the editing of two codons in Region II (G110E+A111T) does not result in any apparent change, a more extensive, three-codon mutation (M109I+G110K+A111T) results in a lack of wild-type level eIF4E1 protein expression due to the altered splicing in the genomic sequence.

### 5. A combination of mutations in Region I and II of *eIF4E1* capacitates resistance to several isolates of potyviruses

Finally, we evaluated the resistance spectrum to potyviruses of all the 10 edited tomato lines—two for each allele, along with plants of wild-type WVa106, TILLING *eif4e1^KO^* mutant, and a *pot1* near-isogenic line (NIL-pot1) (Lebaron et al., 2016). Plants were challenged with three isolates of PVY and two other potyviruses, PepMoV and tobacco etch virus (TEV) CAA10 isolate, previously tested (Gauffier et al., 2016) Resistance of each genotype was based on viral accumulation assessed by ELISA after 21 days.

All eight lines harboring Region II mutations ET or IKT, regardless of the presence of P69T in Region I, showed significant resistance to PepMoV and PVY-LYE90 (Table 2). This is significant but not any better than the limited resistance spectrum associated with knocking out *eIF4E1*. Indeed, the limited resistance spectra of the *eIF4E1^IKT^* and *eIF4E1^P69T/IKT^* mutants were consistent with their loss of function of the *eIF4E1* gene just like in the TILLING *eif4e1^KO^*. In contrast, the combination of mutations P69T and ET in eIF4E1 (eIF4E1^P69T/ET^) triggered significant resistance to PVY-N605, as well as partial resistance to the PVY-SON41 isolate (Table 2, Fig. 5A). The infection of PVY-N605 was further assessed by inoculating a GFP-tagged infectious clone of this isolate and monitoring the GFP fluorescence. GFP fluorescence could be observed on the inoculated and systemic leaves of the wild-type WVa106 and of the *eIF4E1^ET^* line, mutated only in Region II. On the contrary, the fluorescence was observed neither on the inoculated nor the systemic leaves of the *eIF4E1^P69T/ET^* lines and NIL-pot1, thereby confirming the resistance of the specific Region I + II edited tomato (Fig. 3B). Nevertheless, the *eIF4E1^P69T/ET^* allele was not associated with resistance to TEV-CAA10 unlike the natural resistance *pot1* allele in which a larger set of amino acid substitutions has been selected (Table 2).

**Table 2.** Schematic diagram of the selected edited alleles.

| Genotypes | Region I | Region II |  | Virus strains tested |  |  |  |  |
| --- | --- | --- | --- | --- | --- | --- | --- | --- |
|  | P69T | G110E+ A111T | M109I+ G110K+ A111T | PepMoV | PVY-LYE90 | PVY-N605 | PVY-SON41 | TEV-CAA10 |
| WVa106 (wildtype) |  |  |  | S | S | S | S | S |
| TILLING *elf4e1<sup>KO</sup>* |  |  |  | R | PR | S | S | S |
| NIL-pot1 |  |  |  | R | R | R | R | R |
| P69T | X |  |  | S | S | S | S | S |
| ET |  | X |  | R | PR | S | S | S |
| P69T/ET | X | X |  | R | R | R | PR | S |
| IKT |  |  | X | R | PR | S | S | S |
| P69T/IKT | X |  | X | R | PR | S | S | S |

Summary of virus infection testing results of each mutant against five potyvirus species. The accumulation of viral coat protein (CP) after 21 dpi was measured by ELISA. Six to ten plants were used for each mutant line and 12-15 plants for wildtype. Two independent lines were tested for the five edited genotypes. S: susceptible, R: resistant (colored in yellow), PR: partially resistant (CP level lower than S but higher than R; colored in orange).

**Figure 5.**
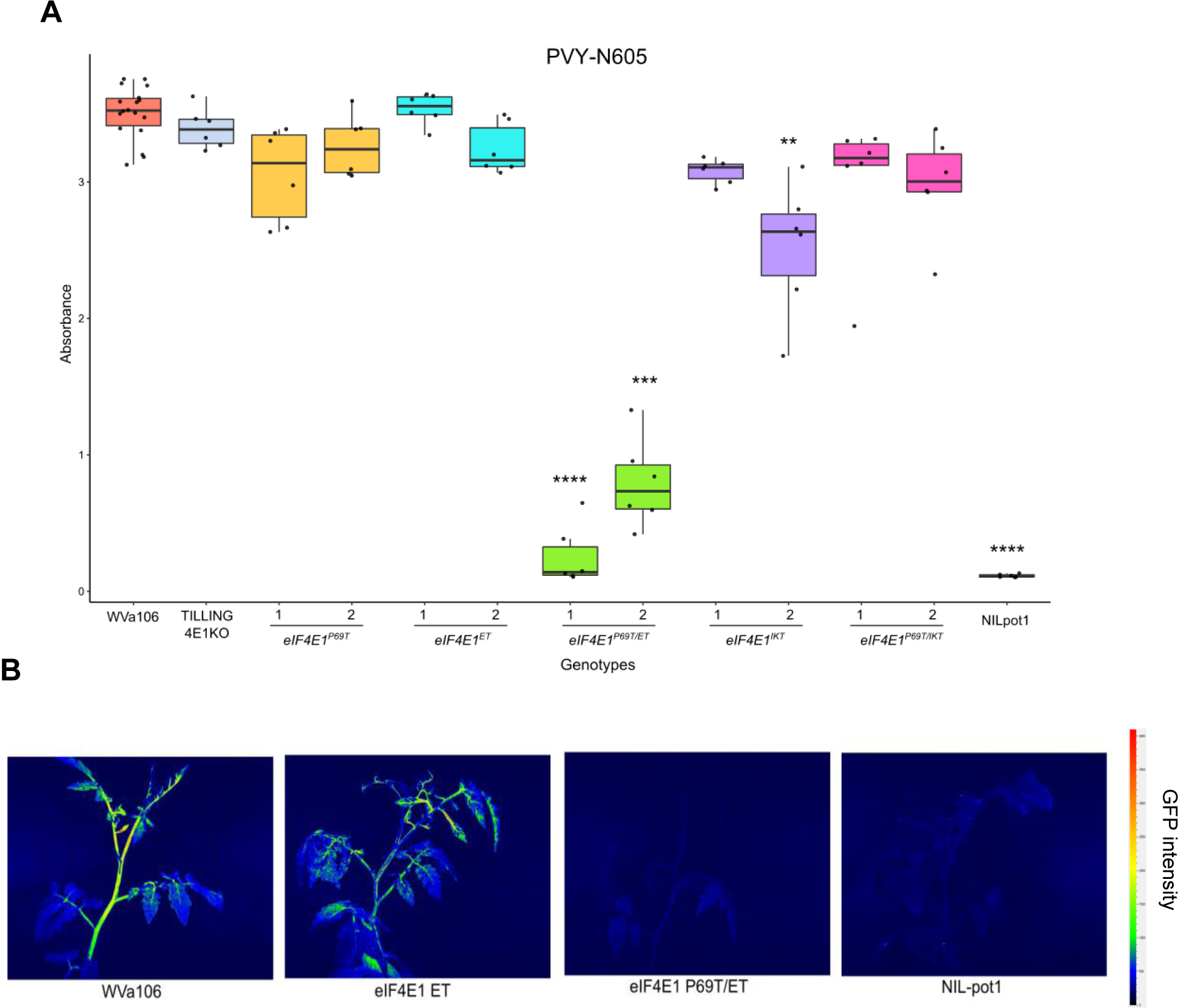
A combination of mutations at eIF4E1 region I and II triggers resistance to PVY N605. **A** DAS-ELISA quantification of the accumulation of PVY-N605 in tomato plants at 21 dpi. Points overlaid on boxplots represent the individual plants. Wild-type WVa106 and KO4E1 (TILLING *eif4e1^KO^*) lines were used as positive controls, while NIL-pot1 plants are used as controls for resistance. Two independent lines were tested for the five edited genotypes. Colored by alleles. Six plants were tested per mutant and 12 for wildtype. Significance of difference compared to WVa106 based on Kruskal-Wallis and Dunn-test as a *post-hoc* test is indicated by asterisks. < 10^-4^ ****; 10^-4^ to 10^-3^ ***; 10^-3^ to 10^-2^ **. **B** Monitoring of the virus infection using a GFP-tagged PVY-N605 clone corroborates the resistance of the edited *eIF4E1^P69T/ET^* line demonstrated by ELISA results. Wild-type WVa106 was used as the positive control, while NIL-pot1 was used as the resistant control. GFP signal was observed at 15 dpi under blue light (420 nm) by using GFP camera imaging. The bar next to the photos indicate the intensity of GFP perceived by the device, from blue (no GFP) to red (high intensity). Each photo is a representative of 10-15 plants observed for each genotype. Two independent lines were tested for the edited alleles.

## Discussion

In this work, we have used TALEN-based gene knock-in and CRISPR-mediated base editing techniques on the tomato *eIF4E1* allele in an attempt to generate genetic resistance against a broad range of potyviruses. Our targeted approach to the specific regions in *eIF4E1* for multiple base substitutions has brought about a *de novo* functional resistant tomato *eIF4E1* allele. Edited plants with this allele showed the broadest potyvirus resistance spectrum achieved by genetic engineering in tomato so far. Meanwhile, we have observed a previously unreported effect on RNA splicing by certain base modifications in a target exon resulting in abnormal expression.

Since the identification of pepper PVY-resistance alleles *pvr1*/*pvr2* as alleles of the *eIF4E1* gene in early 2000s, there has been a number of potyvirus resistant *eIF4E* alleles found among genetic resources, including *pot1* in tomato (Ruffel et al., 2002; Kang et al., 2005; Ruffel et al., 2006). Apart from using transgenic expression, direct implementation of these genes to other species in resistance development was restricted because they are found in a limited number of species and the conventional breeding is often bound by the interspecies barriers. Moreover, the functional significance and redundancy of the *eIF4E* genes in plants make the gene KO or knock-down strategies insufficient (Piron et al., 2010; Mazier et al., 2011; Gauffier et al., 2016; Chandrasekaran et al., 2016; Kuroiwa et al., 2022; Pechar et al., 2022). These results suggest that an alternate strategy, based on the generation of resistant functional alleles, similar to the ones selected among the natural variability, should be more efficient. Thus, more delicate engineering at the base level is clearly needed to achieve resistance and keep eIF4E function at the same time (Bastet et al., 2017).

Although this work confirms the high efficiency of base editing to generate diversity in a specific region of a gene, we also highlight a potential drawback of base editing following the mutation of four bases in the *eIF4E1* Region II. While the genomic modification in the IKT- coding mutants was done correctly, the expression of the resulting full-length mRNA and protein was impaired because the genomic modification induced an exon skipping. So far, there are no reported cases of such aberrant splicing due to CRISPR-mediated editing in exons of a gene although there are some examples after editing in introns or inducing indels in exons (Li et al., 2019; Tang et al., 2021). It is possible that this set of modifications has converted the exon 2 into an intron-like sequence. Introns are often defined by UA-richness of the sequence (Simpson et al., 1996). The modified sequence does not completely correspond to known consensus of branchpoint or intronic splicing enhancer sequences which are important intronic *cis*-elements in splicing events (Wang et al., 2012; Zhang et al., 2019). Nevertheless, it is tempting to associate the increased A-richness in an otherwise exonic region, with the false recognition of the intron. On a practical point, our results stress that the sequencing of the genomic target is not sufficient for inferring the effect of a base editing strategy on a gene. This holds an important message that i) the results of the genome-editing should be confirmed by assessing cDNA and protein expressions and that ii) codon choices should not be neglected.

Here, by inducing multiple amino acid substitutions in eIF4E1, we have designed cherry tomato mutant lines which present resistance against a wider range of potyviruses than the single *eif4e1^KO^* mutant. To our knowledge, this is the first example of viral resistance conferred to a crop species via base-editing technology by generating a *de novo* resistance *eIF4E1* allele. The amino acid changes induced in the resistance eIF4E1 include side-chain-lacking Glycine to negatively charged Glutamic Acid at position 110 (G110E). Since potyviral VPg is negatively charged, such substitution may repel the VPg protein from the two Regions and reduce the interaction. Nevertheless, this substitution does not abolish the functionality of the resistant protein as shown in the cap-affinity purification and yeast complementation assay. Remarkably, tomatoes possessing mutant eIF4E1^P69T/ET^ have accumulated a very low level of PVY-N605 which is known to use either *Sl*eIF4E1 or *Sl*eIF4E2 in the absence of one of them in susceptible wild-type cultivars (Gauffier et al., 2016; this work). The resistance against this strain had been achieved only when the two eIF4Es were knocked-out or when the *SheIF4E1-pot1* allele from the wild relative *S. habrochaites*, was expressed (Gauffier et al., 2016; Lebaron et al., 2016). Our results show that with the exception of the TEV-CAA10, the eIF4E1^P69T/ET^ protein behaves similarly to the natural resistant *pot1*-encoded protein that appears to make the *Sl*eIF4E2 unavailable to the potyvirus infection by a yet unknown mechanism.

In all, our deliverables prove that it is possible to pyramid modifications in two separate regions in one gene to establish resistance while sustaining the functionality and that indeed a combination of mutations in the two Regions is necessary to establish resistance. To induce targeted genetic variation in the tomato *eIF4E1*, we have focused on two separate domains in the genome that are in the proximity in the predicted 3D protein structure (Supplemental Fig. 5) and achieved virus resistance.

Our work summarizes the progress of base editing in the recent years: from gene knock-in using TALENs to the CRISPR-based base editors. In the coming years, we expect that a similar strategy could be designed and carried on in a faster and more accurate manner, by using prime editor and/or CRISPR/Cas proteins with relaxed PAM requirement (Veillet et al., 2020; Ren et al., 2021). These techniques should allow us to tinker the protein function as we design and to minimize the unfavorable effects brought about by the insertions/deletions or lack of precision. By taking the susceptibility factor eIF4E as an example, we exemplify how genome editing can be used to design complex alleles and generate useful traits that could boost plant breeding.

## Material and methods

### Plant material

Cherry tomato (*Solanum lycopersicum* var. *cerasiforme*) cv. West Virginia 106 (WVa106), was used as background for the editing of *eIF4E1* (Solyc03g005870) and as the wildtype. The TILLING line *eif4e1^KO^*was previously obtained through ethyl methanesulfonate (EMS) mutagenesis in the *S. lycopersicum* cv. M82 background (Piron et al., 2010). NIL-pot1 is a near-isogenic line of the *eIF4E1* allele *pot1* from *S. habrochaites* PI247087 introgressed in *S. lycopersicum* cv. Mospomorist (Lebaron et al., 2016). All plants were grown in a growth chamber at 24°C/18°C in the 16 hr day/8 hr night cycle.

### Constructs for genome editing using TALEN^®^, CRISPR/Cas9, and Cytosine Base Editors (CBE)

TALENs^®^ designed to target 5′- TTTCCTCTTTCAAATTggtgatagtgtagtgTAAGGGAAAACAGGGA (nucleotides recognized by the TAL effectors are capitalized) in the *SleIF4E1* genomic sequence were cloned into vectors, pCLS23222 and pCLS23224, and provided by Cellectis (Paris, France) (Supplemental Data 2, 3). The DNA donor template for gene knock-in was synthesized by GenScript (Piscataway, NJ, USA) and cloned into pBGWD0 vector (Gateway vector VIB) through LR Gateway recombination. The linear DNA template designed for HR was generated by digestion with PacI and PmeI (Fig. 1B, Supplemental Data 4). For the two targeted deletion steps using CRISPR/Cas9 after the NHEJ event, different pairs of single guide RNAs: pair 1 [sgRNA- DKM04 and sgRNA-eIF4E05] and pair 2 [sgRNA-Cl-01 and sgRNA- Cl-02] were cloned into pDe-Cas9-Hpt binary vector and pDe-Cas9-NptII binary vector, respectively (Danilo et al. 2018). The sgRNA cassettes were synthesized by Integrated DNA Technologies (Coralville, IA, USA). For co-base-editing of the *SleIF4E1* and *SlALS1*, an *SleIF4E1* sgRNA cassette with *StU6* promoter was cloned into pDicAID_nCas9-PmCDA1_NptII binary vector inserted with *AtU6p*-*SlALS1* guide (Veillet et al., 2019) (Supplemental Fig. 3A and B, Supplemental Data 5) and named CBE-4E1-RII. The *SleIF4E1* sgRNA cassette was synthesized by TwistBioscience (San Francisco, CA, USA). TALEN targets and CRISPR guide RNA sequences are listed in the Supplemental Table 1.

### Genetic transformation of tomato

TALEN^®^ plasmids and the linear DNA template were delivered into leaves from four-week- old tomato plants by biolistic method. Other transformations with CRISPR/Cas9 or CBE vectors were performed on cotyledon pieces from 8-12-day-old tomato seedlings by Agrobacterium-mediated transformation using *Agrobacterium tumefaciens* C58pGV2260 strain. Plant regeneration and transformant selection following both transformations were performed as previously described (Mazier et al., 2011; Danilo et al., 2018; Veillet et al., 2019). Detailed methods are available in supplemental material and method. *In vitro* culture was carried out in a controlled environment at the temperature of 22°C/18°C with 16 hr/8 hr (day/night) photoperiod.

### Genotyping by PCR and High-Resolution Melting analysis

Genomic DNA was extracted from fresh tomato leaf samples according to (Fulton et al., 1995). T0 plantlets after editing with TALEN and CRISPR/Cas9 were screened by PCR using specific primers for each transformation. T0 plantlets after co-base-editing using CBE were screened by HRM analysis using specific primers and Precision Melt Supermix (BioRad, Hercules, USA) with the CFX96™ Real-Time PCR Detection System (BioRad), as previously described (Veillet et al., 2019). Melting-curve analysis was performed using the Precision Melt Analysis software (BioRad, Hercules, USA). PCR was also performed on the *SleIF4E1* genomic sequence to further verify. *GAPDH* (*Glyceraldehyde-3-Phosphate Dehydrogenase*) gene was amplified as the technical control with gapdhF/R. PCR products were Sanger-sequenced by Genoscreen (Lille, France). T-DNA detection was done by PCR. All the primer sequences and pairs used for detection are available in Supplemental Table 2.

### Total RNA extraction and cDNA analysis

Total RNA was extracted from young tomato leaf tissue (< 100 mg) of each plant in 500 µL of TRI-Reagent (Sigma-Aldrich, St. Louis, USA) and quality-checked by concentration measurement on Nanodrop ND-1000 (ThermoFisher, Waltham, USA). Reverse transcription (RT) of the extracted RNA samples (1 µg) was performed using AMV (Avian Myeloblastosis Virus) Reverse Transcription Kit (Invitrogen, Waltham, USA) and oligo-dT18 (1 µM) to obtain coding DNA (cDNA). The cDNA for *eIF4E1* was amplified with a primer pair c4e1F/R and *GAPDH* (*Glyceraldehyde-3-Phosphate Dehydrogenase*) gene as the internal control with gapdhF/R.

### Protein extraction and western blot analysis

Total protein was extracted from 100 mg of young tomato leaves, and 150 µL of Laemmli 2x DTT buffer was added to the crushed leaves. The homogenized solution was boiled for 5 min. The supernatant after centrifugation at 4°C was collected and conserved at -20°C. Western blot was performed as previously described using specific antibodies (Kuroiwa et al., 2022). The anti-*Sl*eIF4E1 polyclonal serum (Gauffier et al., 2016) was diluted 1/2000, and combined with secondary goat horseradish peroxidase-labelled anti-rabbit serum (Sigma-Aldrich, St. Louis, USA) diluted 1/5000. As loading controls, monoclonal anti-plant actin antibodies (1/10000 dilution) (Sigma-Aldrich) were used with horseradish peroxidase-labelled anti-mouse serum (1/10000 dilution) (Sigma-Aldrich). Dilution was done in TBS 1x + 5% milk (Tween 0.05% + Triton X-100 0.2% for cap-affinity purification).

### Virus isolates and infection assays using DAS-ELISA quantification

Potyvirus infection assays were performed using PepMoV (Mazier et al., 2011), PVY-N605 (Parrella et al., 2002), PVY-LYE84, PVY-SON41 (Moury et al., 2004), and TEV-CAA10 (Charron et al., 2008). Isolates were maintained in *Nicotiana benthamiana* plants 12 days before inoculation. For the PVY infection monitoring, the infectious clone of PVY-N605 tagged with GFP (hereafter, PVY-N605-GFP) incorporated in pCambia vector was multiplied on *N. tabacum* cv. Xanthi plants before inoculation of 21-day-old tomato plants. Inocula were prepared from 1 g (fresh weight) of the infected tobacco leaves ground with 4 mL of grinding buffer: potassium phosphate buffer (0.03 M, pH = 7) with 0.2% of diethyldithiocarbamate, 80 mg active charcoal and 80 mg carborundum. Six to twelve plants per genotype were mechanically inoculated on the cotyledons of 14-day-old tomato plants. At 21 days post-inoculation (dpi), 1 g of leaves from each plant was collected and ground in 4 mL of grinding buffer. To measure the virus accumulation, DAS-ELISA was performed using anti-PVY or anti-TEV antiserum (Sediag, Bretenière, France) or anti-polypoty antiserum (Agdia, Grigny, France) for PepMoV according to the manufacturer’s instructions. The absorbance was measured by a spectrophotometer at wavelength 405 nm.

### Virus detection by fluorometric camera analysis

Monitoring of PVY-N605-GFP infection was carried out with a closed fluorometric camera FluorCam FC 800-C/1010- GFP (Photon System Instruments, Drasov, Czech Republic) equipped with a GFP filter. For each genotype, 10-15 plants were used. GFP fluorescence was captured by the camera upon excitation at 420 nm and the intensity of the GFP signal was indicated by the color scale.

### m^7^GTP cap-affinity purification

Total soluble proteins were extracted from 300 mg leaf tissue in 1200 µL binding buffer with protease inhibitor (+PI) (performed in three sets 100 mg:400 µL each). After centrifugation at 13,000 rpm for 10 min at 4 °C, 50 µL of the supernatant (INPUT) was recovered, and the rest was incubated with 150 µL of γ-aminophenyl-m^7^GTP (C10-spacer)-agarose beads (Jena Bioscience, Jena, Germany) pre-equilibrated with binding buffer without protease inhibitor at 4°C for 1.5 h. The beads were pelleted for 1 min at 13,000 rpm and washed three times with binding buffer +PI at 4°C to remove unbound proteins. The INPUT was diluted two-fold with Laemmli DTT 2x buffer. The proteins linked to the cap analog were eluted using 100 µL Laemmli DTT 2x buffer (OUTPUT) and analyzed by western blot assay as described above.

### Yeast complementation assay

Amplified cDNA from the different *eIF4E1* alleles was subcloned using the gateway technology (ThermoFisher) in a converted p424-GDP-DEST plasmids. Yeast complementation method was adapted from (Gauffier et al., 2016). Briefly, attB sequence was attached to cDNA for BP recombination reaction to generate the entry clones. The entry clone was then used for LR recombination reaction with the destination vector p424-GPD to generate the tryptophan- selectable expression vector with a constitutive promoter pGAD. Then, yeast (*Saccharomyces cerevisiae*) strain J055 (cdc33-Δ: LEU2Leu2 ura3 his3 trp1 ade2 [YCp33supex-h4E URA3]) that lacks its native SceIF4E and expresses human eIF4E (H eIF4E) under galactose-inducible, glucose-repressible promoter pGAL was transformed with each expression vector. The transformants were selected on galactose-containing, uracil/tryptophan deficient media (-UW); then, the transformed colonies were spotted on glucose-containing, tryptophan deficient media (-W) to see if the eIF4E expressed from the expression vector complements the lack of its native eIF4E or repressed H eIF4E. Incubation for the complementation assay was done at 20°C/16°C with 16 hr/8 hr cycle. The empty vector is p424-RfA.

### 3D modelization of eIF4E structures

Homology modelling of the eIF4E2 protein were carried out as previously (Moury et al., 2020) using the YASARA software (http://www.yasara.org/), using structural data from pea (Pisum sativum) eIF4E (GenBank ID: AY423375, PDB ID: 2WMC-C) as the template. Protein structure was visualized using PyMol software (https://pymol.org/).

### Statistical analysis

All the statistical tests were performed on the free software R (https://www.R-project.org/). Kruskal-Wallis non-parametric tests were performed to evaluate the significance of differences between genotypes using the “pgirmess” package. As the *post-hoc* test, Dunn test with p-value adjustment by Benjamini-Yekutieli method or Benjamini-Krieger-Yekutieli method was performed using “dunn.test” package to determine the significance.

## Funding

This work was supported by the French national Research Agency (ANR11-BTBR-0001- GENIUS). The IJPB benefits from the support of Saclay Plant Sciences-SPS (ANR-17-EUR- 0007).

## Author contributions

KK, FV, FN, MM, JLG designed the experiments. KK, BD, LP, CT, FV, FD, MM carried out the experiments. FD, PD contributed new reagents. KK and JLG wrote the manuscript with contributions from all authors. All authors read and approved the final manuscript.

## Acknowledgments

The authors thank Emmanuel Botton (INRAE GAFL) for taking care of tomato plants, Nathalie Truglio (INRAE Pathologie Vegetale) for supplying tobacco plants, Pierre Barret, Caroline Tassy, and Anne Partier (INRAE GDEC) for helpful advices to set up biolistic experiments, and Luc Mathis (Cellectis) for TALEN^®^s design and production.

## Competing interests

TALEN® is a registered trademark owned by Cellectis and a patented technology.

## Supplemental Information

**Supplemental Table 1:**
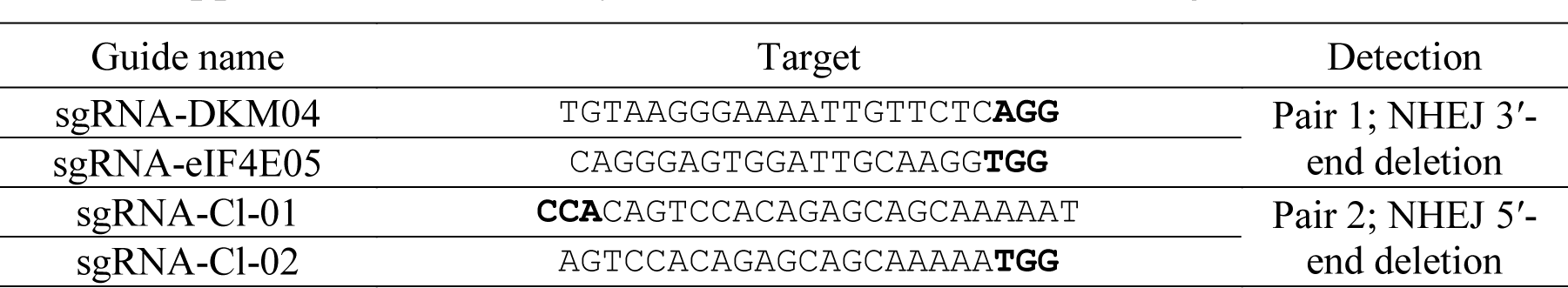
Synthesis of all nucleases and target sites used

**Supplemental Table 2:**
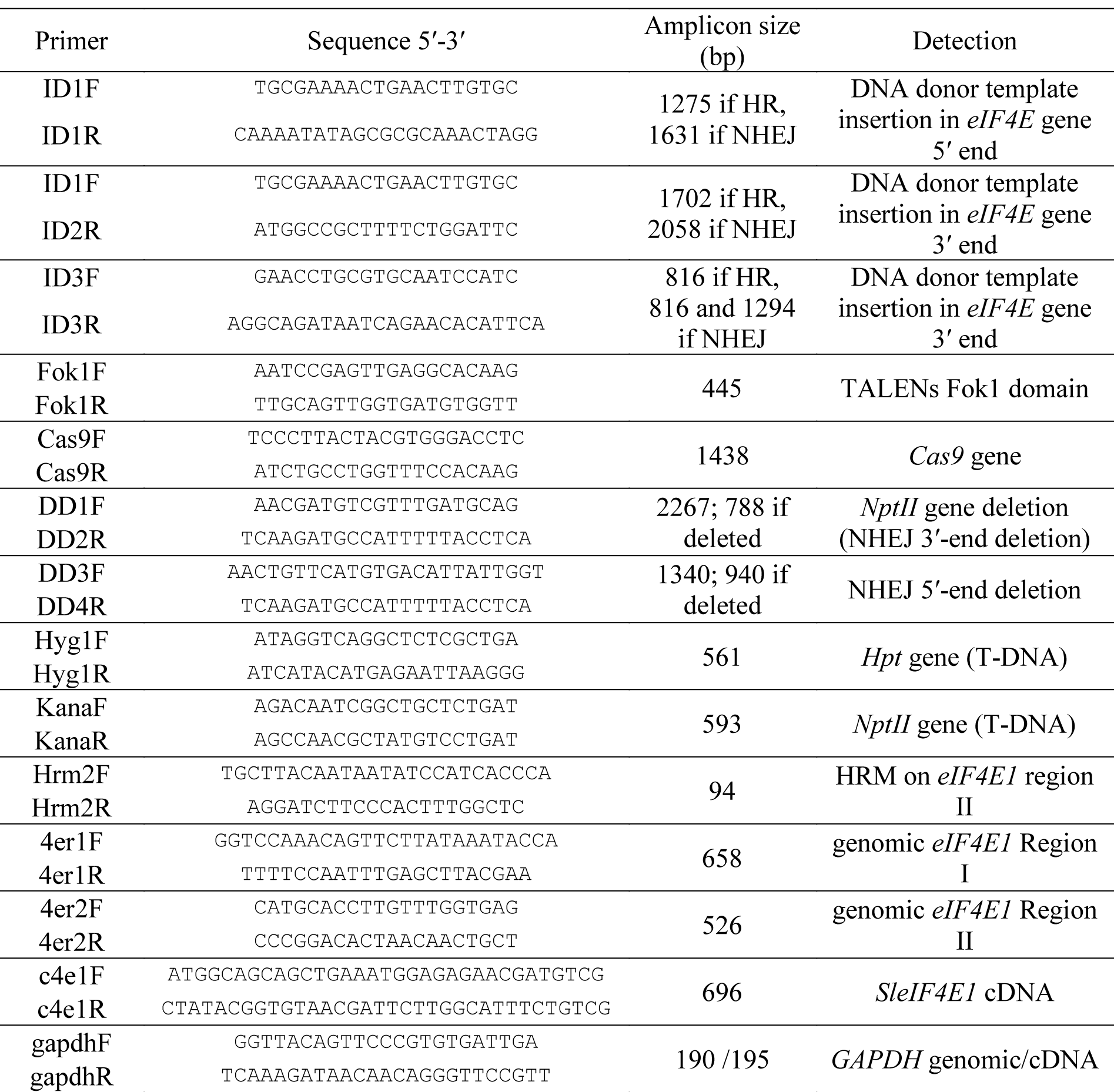
List of primers

**Supplemental Data 1.**
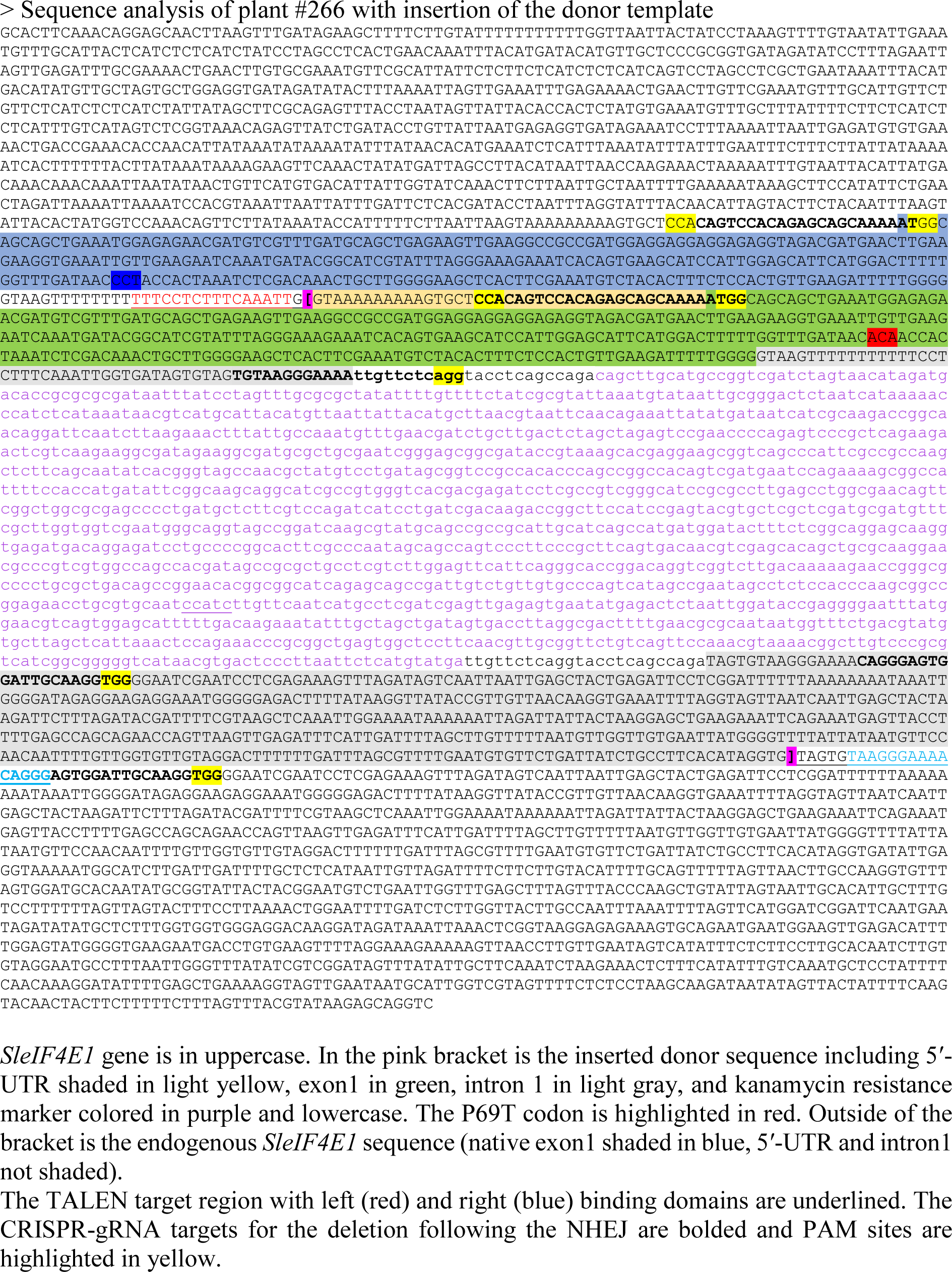

**Supplemental Data 2.**
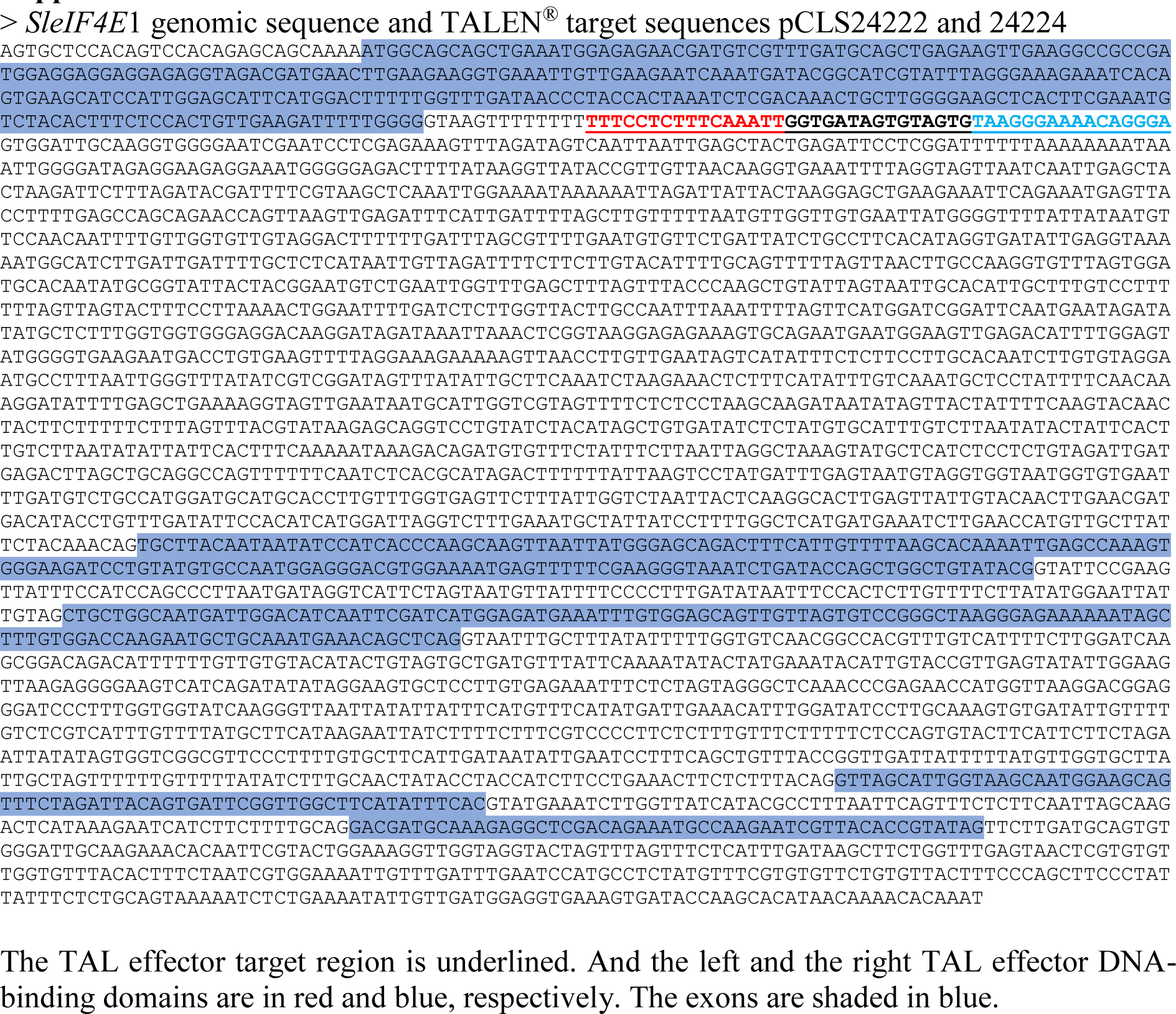

**Supplemental Data 3.**
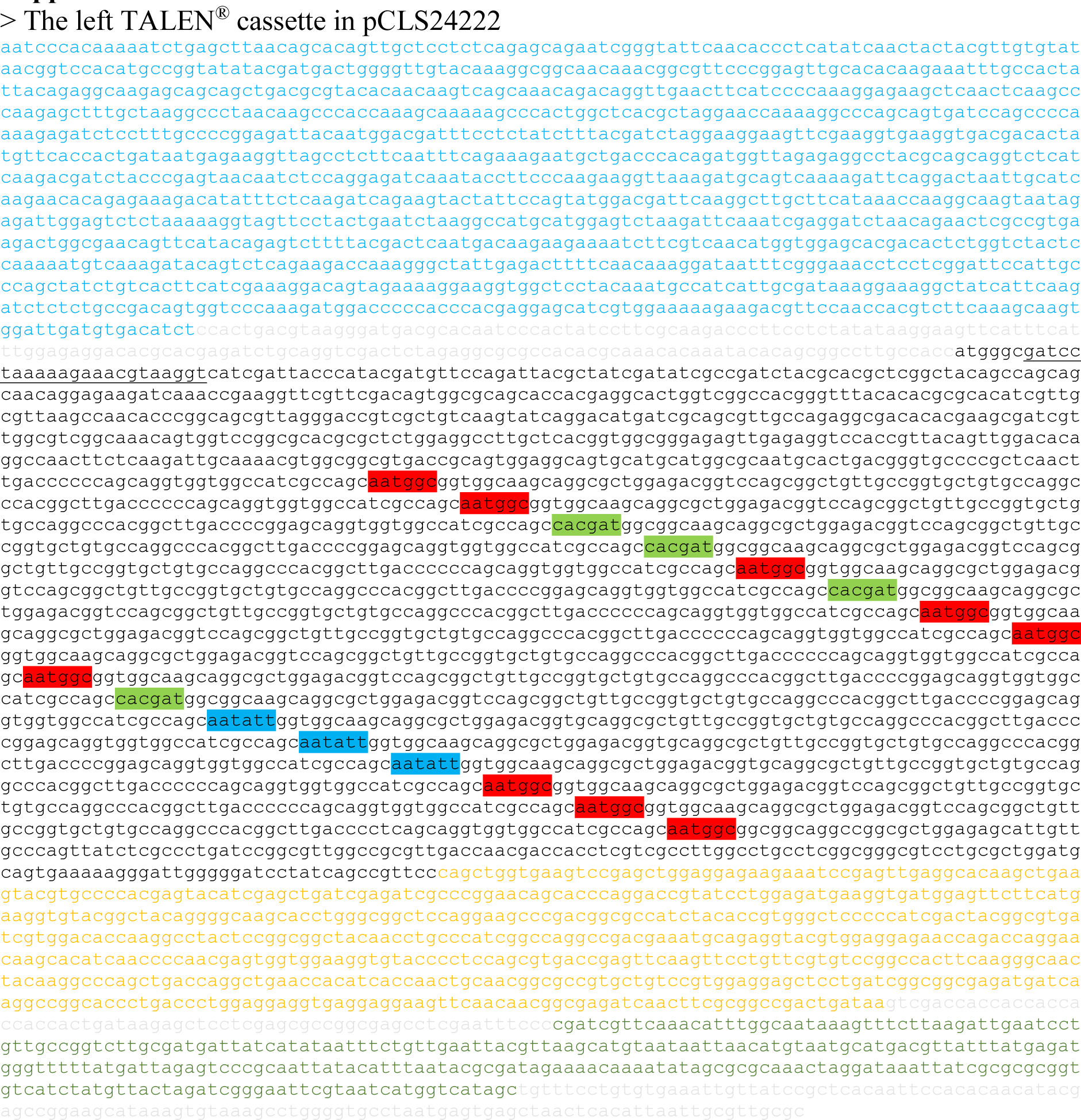
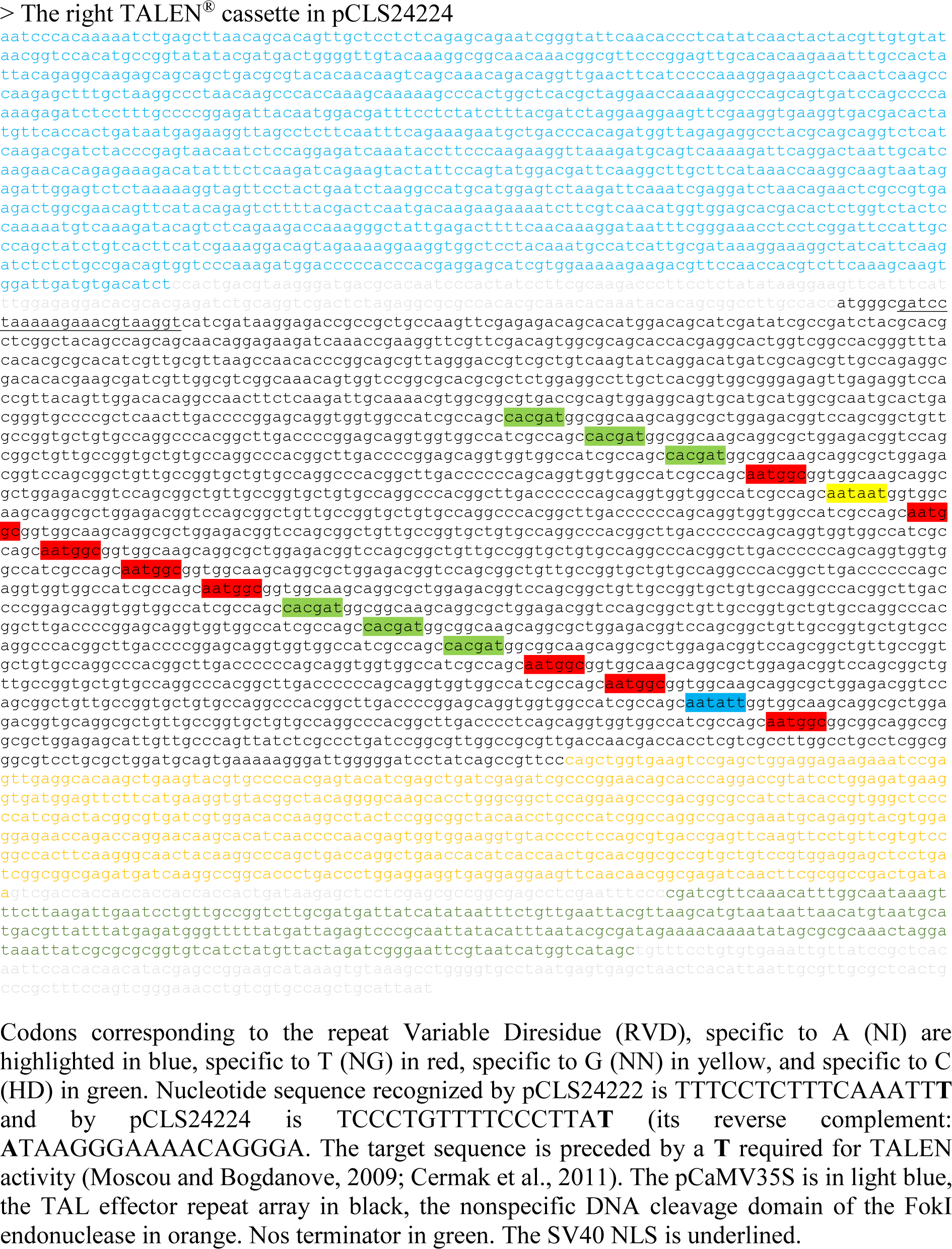

**Supplemental Data 4.**
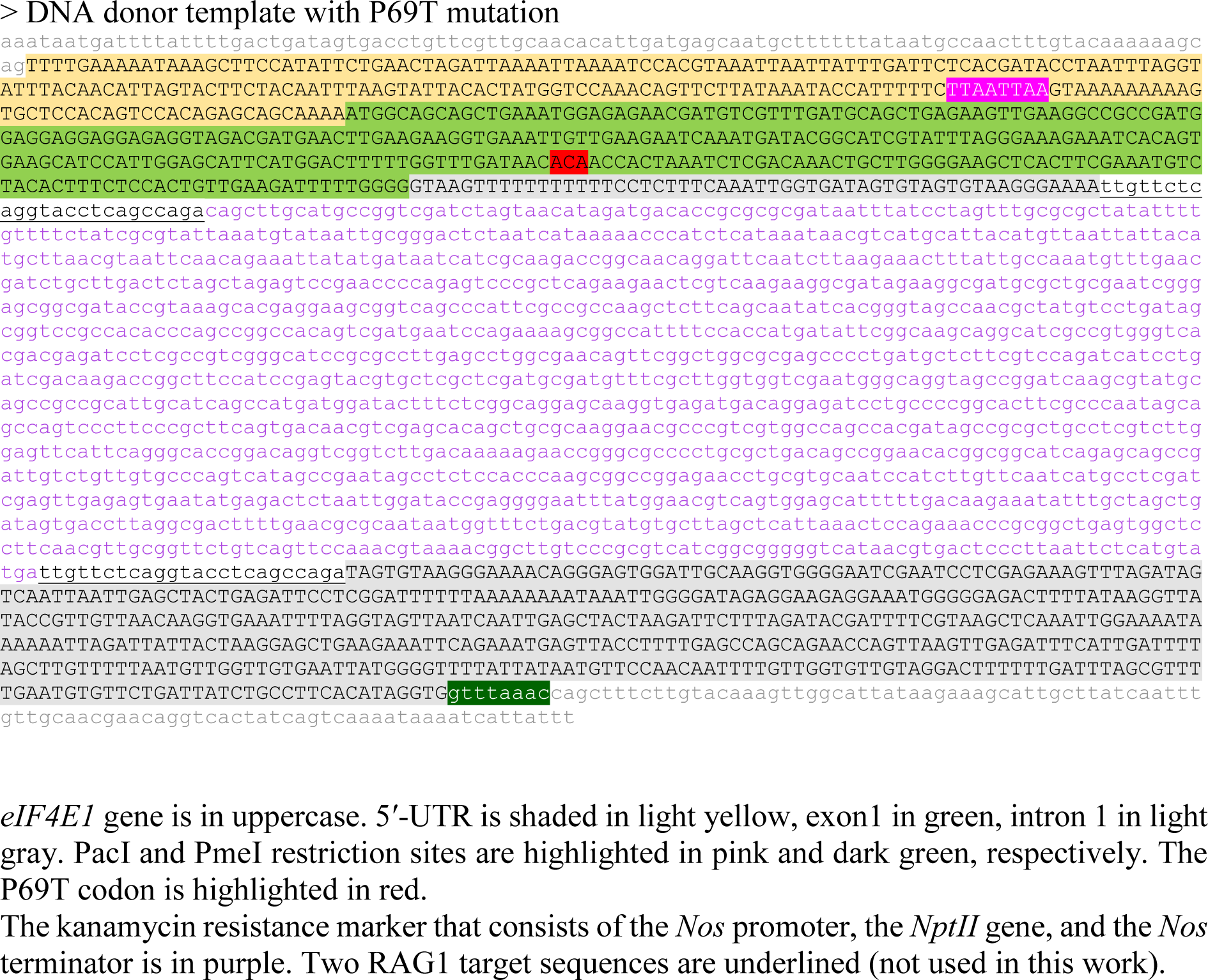

**Supplemental Data 5.**
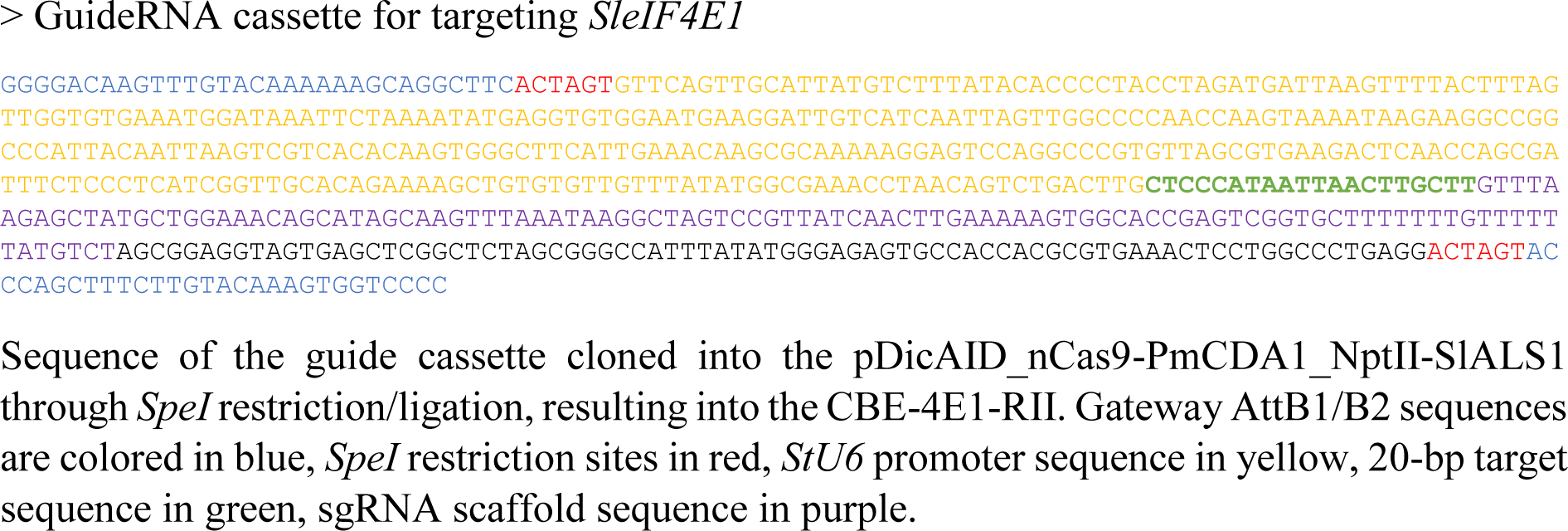

## Supplemental figure legends

**Supplemental Figure 1:**
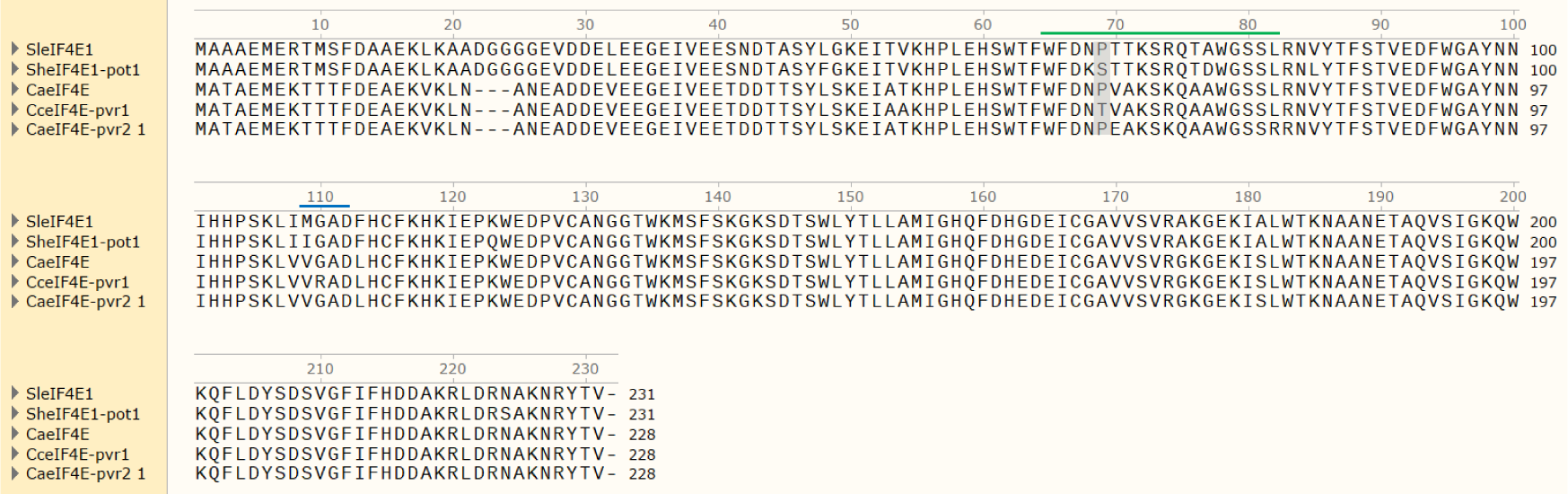
Alignment of the amino acid sequences from potyvirus susceptible and resistant tomato and pepper accessions. “SleIF4E1” and “CaeIF4E” are the representative *eIF4E1* sequence from the susceptible cultivars. “pot1”, “pvr1”, and “pvr1 2” (pvr2^1^) are resistance allele sequences. Green and blue bars indicate the position of the Region I and II, respectively. Tomato P69T and the equivalent pepper P66T are shaded in gray.

**Supplemental Figure 2:**
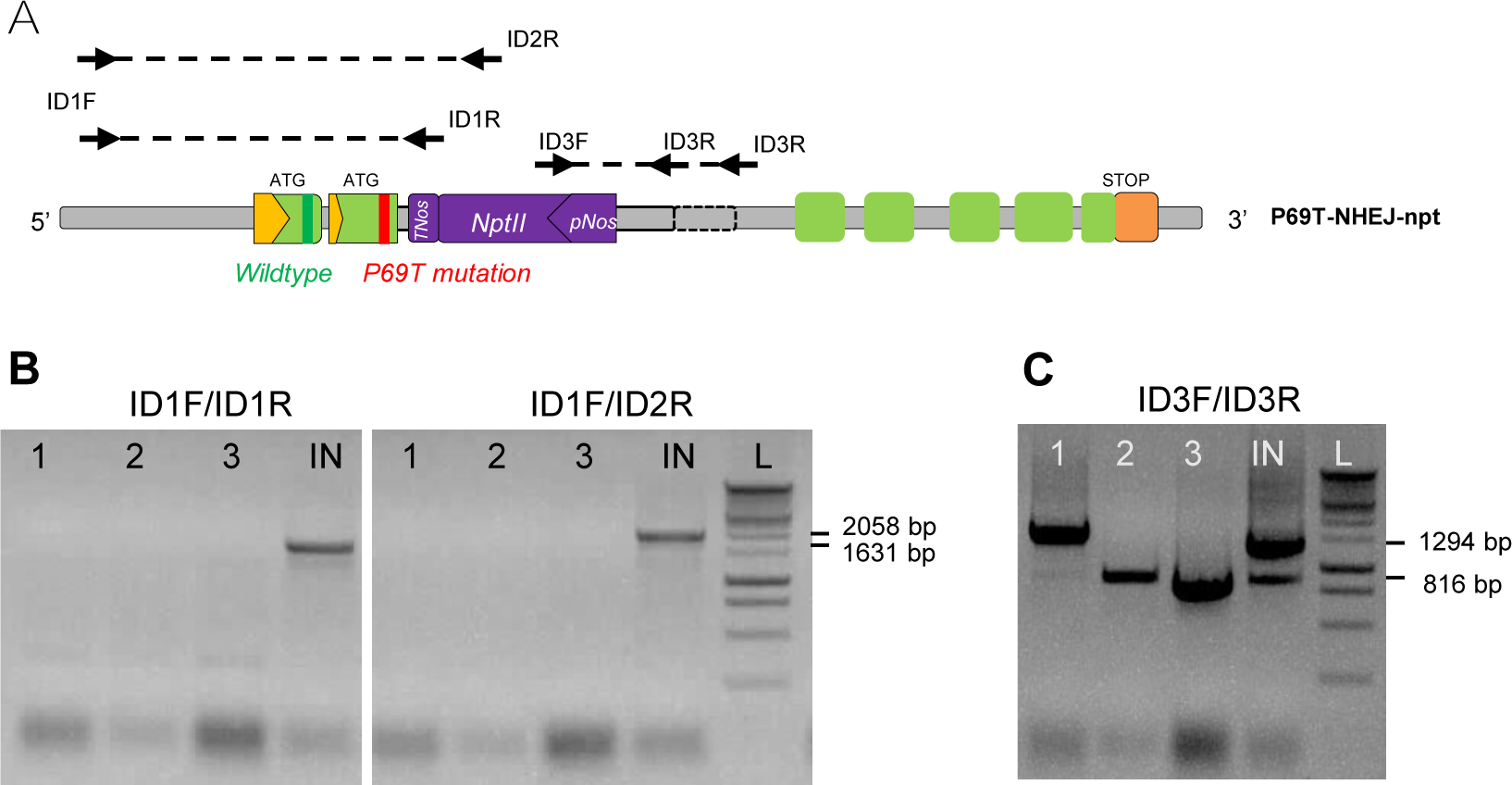
A Primers designed to detect the insertion of the DNA donor template after the biolistic experiment with TALEN. B Detection of the 5′-end of the insert. C Detection of the 3′-end of the insert. The indicated base pairs correspond to the expected band size in an NHEJ event. 1-3 are plant lines with incomplete insertion on the 5′-end. (In C, 1 shows an insertion by NHEJ and 2 and 3 by HR.) IN: NHEJ inserted line (P69T-NHEJ-npt line), L: ladder.

**Supplemental Figure 3:**
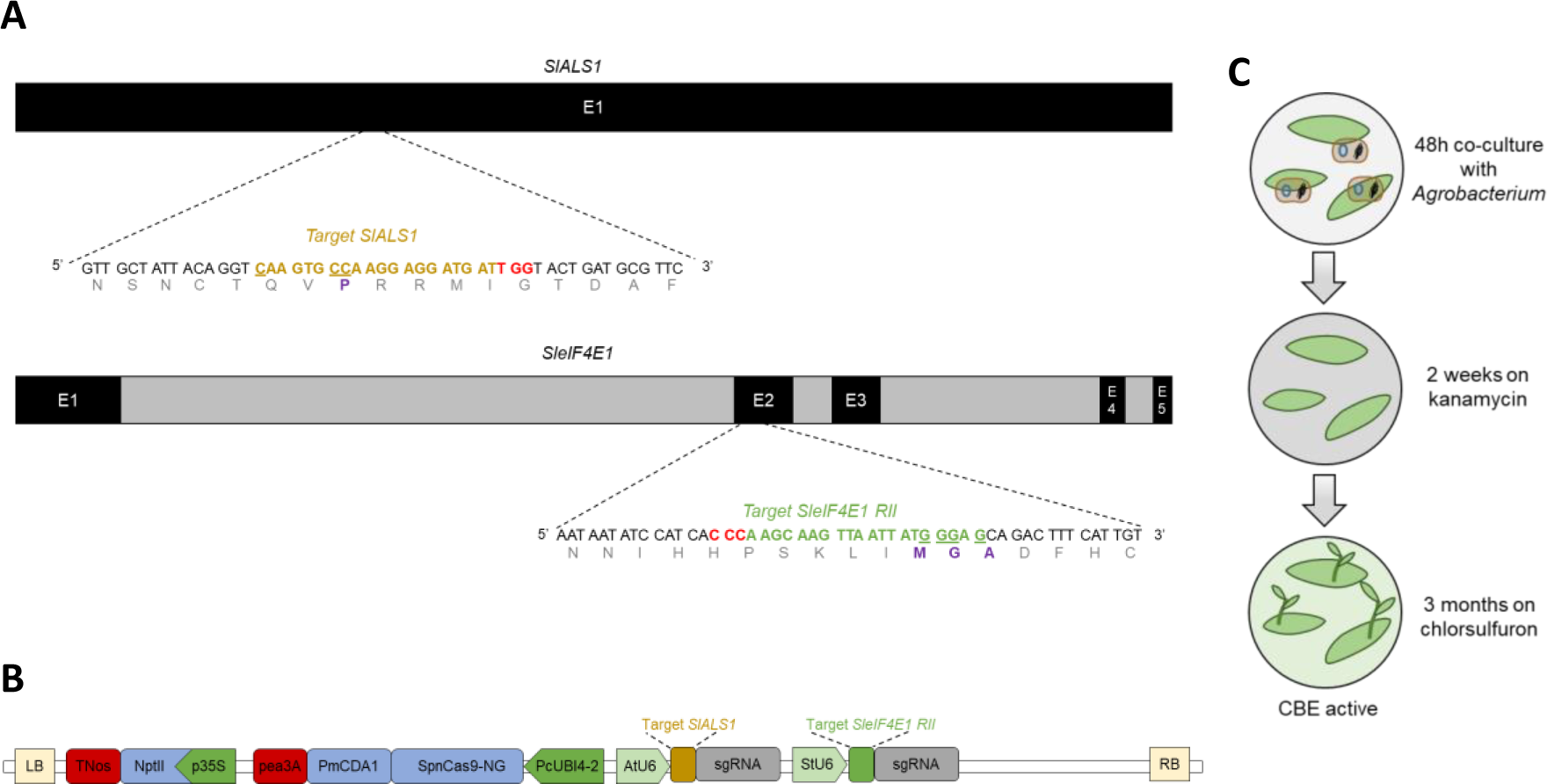
A Target sequences located in the *SlALS1* and *SleIF4E1* genes. Dark and gray boxes represent exons and introns, respectively. The PAM sequence is indicated in red. Targeted nucleotides are underlined and the resulting amino acids are indicated in purple. **B** T-DNA of the CBE-4E1-RII construct used to edit *SlALS1* and *SleIF4E1* genes. LB: left border; RB: right border. **C** Schematic of the production of ALS-co-edited tomato plants through *Agrobacterium*-mediated transformation.

**Supplemental Figure 4:**
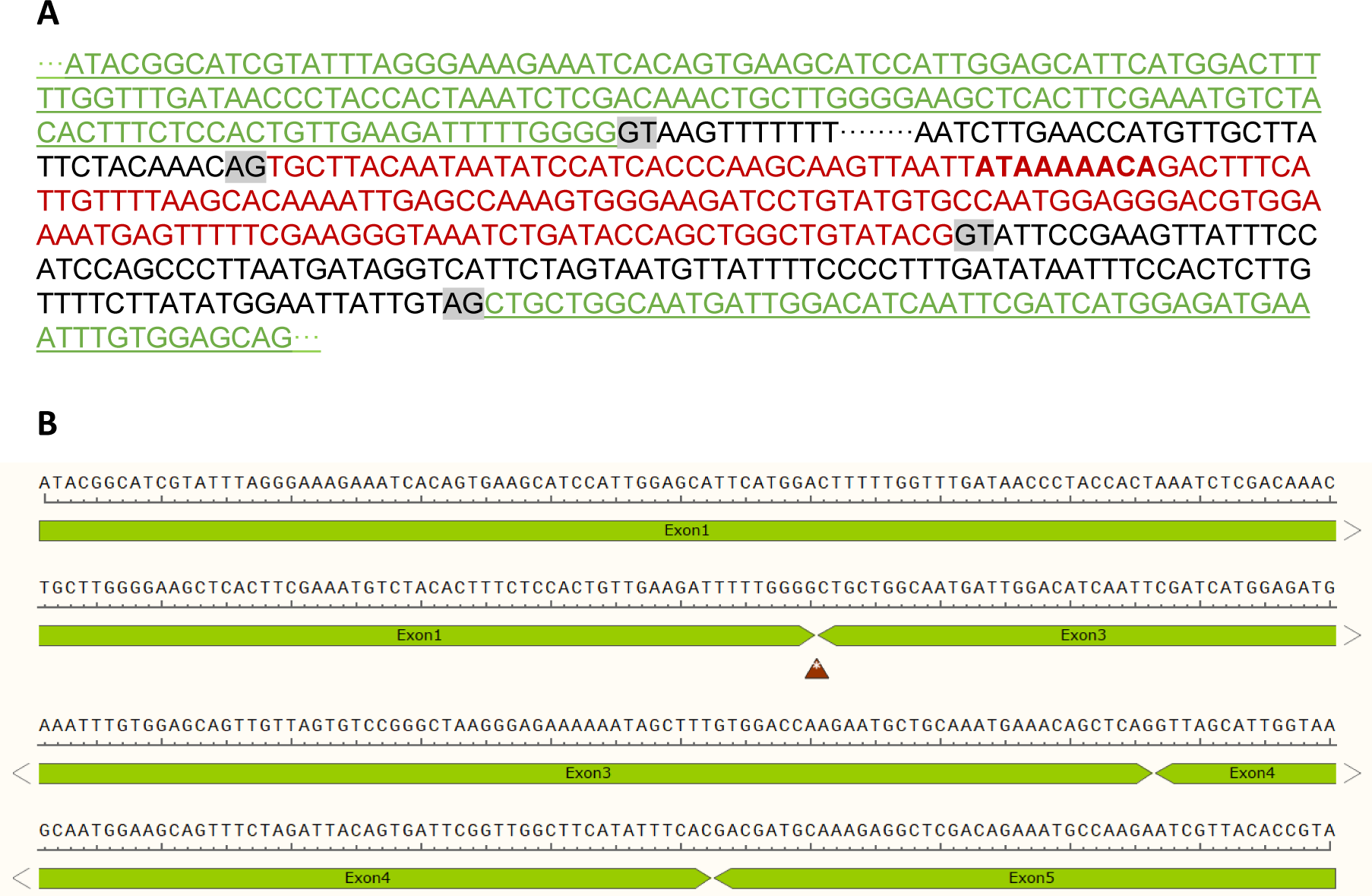
A Genomic sequencing of the Region I and II periphery from an IKT-modified plant line (*eIF4E1^IKT^* line). Shaded in gray are the known splicing donor and acceptor sites. Colored sequences correspond to the exons: exon 1 and 3 in green and exon 2 in red. Underlined sequence is identified in the cDNA sequencing. Bolded is the IKT-coding sequence. **B** A part of cDNA sequenced from the same IKT- modified plant line. Red triangle indicates the site where the wild-type exon 2 is normally found.

**Supplemental Figure 5:**
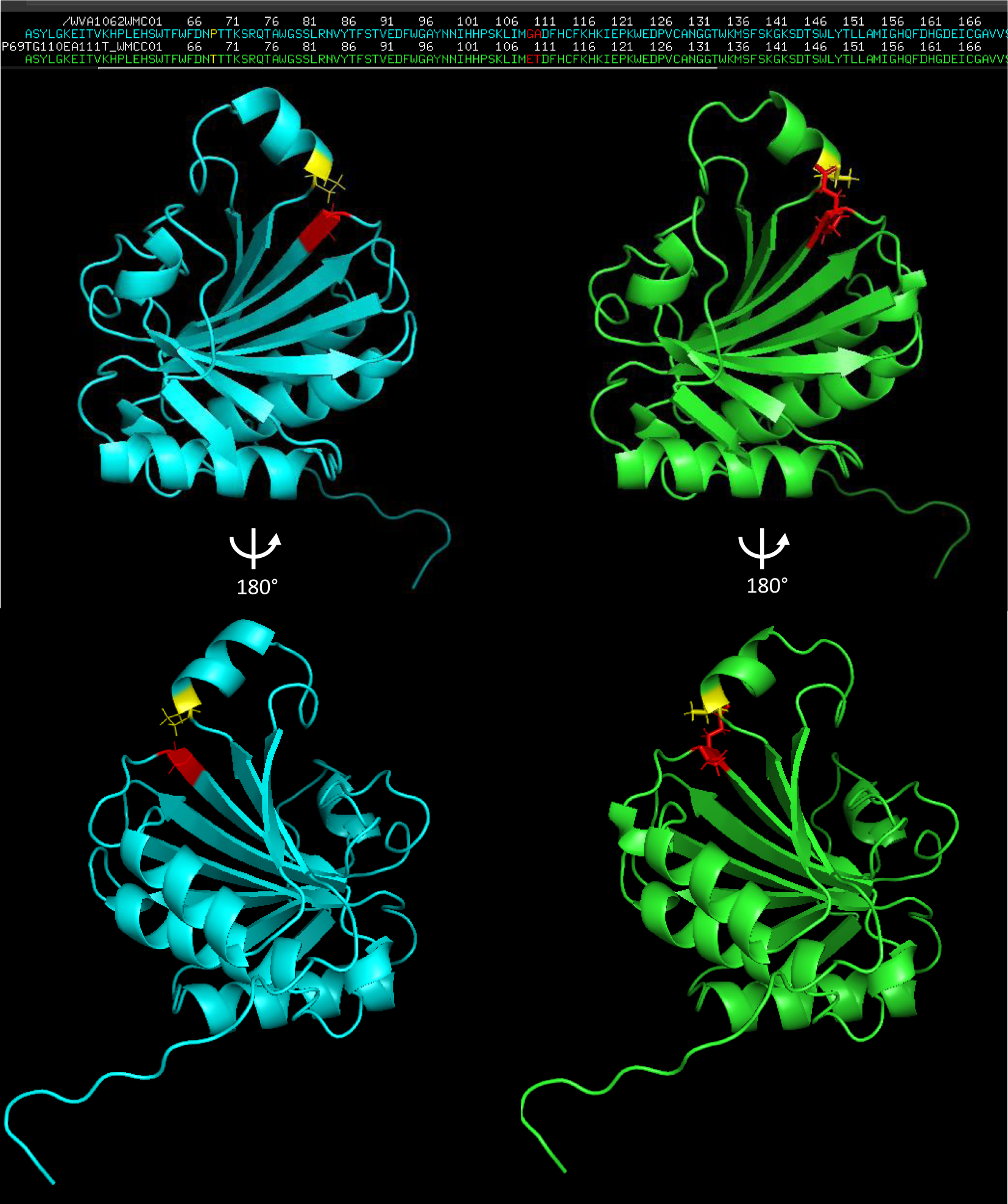
Three-dimensional homology modelling of the tomato eIF4E1 protein, based on crystallography data from the *Pisum sativum* eIF4E 3D structure (PDB ID: 2WMC-C), for the wild-type Wva106 (WT, in blue) and the edited P69T /ET allele (in green). P69, respectively T69 is represented in yellow and G110 A111, E110 T111, respectively in red, with apparent side chains showing. 3D modelization was performed using Yasara and shown with Pymol as in Bastet et al., 2019. 180° rotation along the y axis is represented below.

## Supplemental experimental procedures

### Conditions of genetic transformation via biolistic method

40 leaves from four-weeks-old plants cultivated *in vitro* were used. Leaves were placed on petri dishes containing plasmolysis medium (10% maltose, 0.9 mg/L thiamine, 0.2 mg/L 2-4D, 0.1 mg/L kinetin) for two hours in dark before the biolistic experiments using BioRad PDS-1000/He™ device (Kikkert, 1993) with 1100 psi rupture discs at a target distance of 7 cm. Gold microprojectiles (1 mg) of 0.6 µm diameter were sonicated for 1 min and suspended in 10 µl of DNA solution with spermidine and CaCl2 as described in (Tassy et al., 2014). After the bombardment, leaves were maintained in the same petri dishes during 12 h in dark before they were cut in small 5 mm^2^ pieces and placed on the same medium without maltose for two days in dark.

### Regeneration after genetic transformations

After transformation, leaf pieces were transferred to be selected on the regeneration media containing 2 mg/L zeatin supplemented with appropriate antibiotics (5 mg/L hygromycin or 100 mg/L kanamycin) (Mazier et al., 2011; Danilo et al., 2018). For the co-base-editing, after 10-day incubation on kanamycin media, cotyledon pieces were transferred to fresh selective media containing 40 ng/L of chlorsulfuron every two weeks (Supplemental Fig. 3C) (Veillet et al., 2019). Each regenerating bud was transferred to the culture tubes with selective ½ MS salt elongation medium before genomic screening (Mazier et al., 2011). After the genetic screening, *in vitro* regenerated and rooted T0 plantlets were transferred to greenhouses. T1, T2 and T3 transgenic lines were obtained by self-pollination of the T0, T1 and T2 plantlets respectively. Plants were grown in a sterilized peat soil mixture under natural light conditions.

